# Proteins with amino acid repeats constitute rapidly evolvable and human-specific essentialome

**DOI:** 10.1101/2022.12.29.521938

**Authors:** Anjali Kumari Singh, Ishita Amar, Harikrishnan Ramadasan, Keertana Sai Kappagantula, Sreenivas Chavali

## Abstract

Protein products of essential genes, indispensable for organismal survival, are highly conserved and bring about fundamental functions. Interestingly, proteins that contain amino acid homorepeats that tend to evolve rapidly are enriched in eukaryotic essentialomes. Why are proteins with hypermutable homorepeats enriched in conserved and functionally vital essential proteins? We solve this function versus evolutionary paradox by demonstrating that human essential proteins with homorepeats bring about cross-talk across biological processes through high interactability and have distinct regulatory functions affecting expansive global regulation. Importantly, essential proteins with homorepeats rapidly diverge with the amino acid substitutions frequently affecting functional sites, likely facilitating rapid adaptability. Strikingly, essential proteins with homorepeats influence human-specific embryonic and brain development, implying that the presence of homorepeats could contribute to the emergence of human-specific processes. Thus, we propose that homorepeat containing essential proteins affecting species-specific traits can be potential intervention targets across pathologies including cancers and neurological disorders.

## Introduction

Genome-wide loss-of-function screens have identified essential genes whose deficit results in compromised cell viability or considerable loss of fitness. In unicellular organisms, essential genes characterize organismal essentiality. However, in multicellular organisms, essential genes are often confined to cellular essentiality. This is because gene essentiality is predominantly determined by the cell-type, experimental approaches and culture conditions (*1*). Studies employing variable approaches and culture conditions have identified conditionally essential genes (*2*). In humans, akin to most multicellular organisms, both cellular essentiality and conditional essentiality would contribute to organismal essentiality. Essential genes in human cell lines have been identified using CRISPR-Cas9 mediated genome-wide mutation screens, insertional mutagenesis, gene-trap method and RNA interference (*1–11*). From a functional standpoint, essential genes have been shown to be involved in core biological processes such as gene-expression regulation, signaling, metabolism and/or development in diverse organisms ranging from budding yeast to *Caenorhabditis elegans* to mouse and humans (*7*). Owing to their functional importance, essential genes, compared to non-essential genes, show higher sequence conservation and tend to evolve slowly, predominantly guided by purifying selection (*12, 13*).

Abnormally long repeats of identical amino acids (referred as homorepeats; HRs), such as polyglutamine and polyalanine in proteins lead to detrimental outcomes such as neurological disorders (*14, 15*). Though HRs were conventionally associated with deleterious effects, molecular and systems level studies by others and us identified that HRs play different functional roles such as facilitating molecular interactions, sub-cellular localization, regulating functions and assembly of phase-separated condensates and contribute to conferring adaptability to changing environments (*16–21*). From an evolutionary standpoint, amino acid homorepeats are hypermutable elements that aid in rapid accumulation of genetic variation (*17, 22–24*). Strikingly, *Saccharomyces cerevisiae* essentialome is enriched for proteins that contain homorepeats (HRs) (*17*). These findings present a functional versus evolutionary paradox as to why are these hypermutable elements (homorepeats) prevalent in essential proteins with conserved functionalities. In this study, we analyze the human proteome and present a comprehensive analysis to address this conundrum and elucidate the functional roles and evolutionary benefits associated with homorepeat containing human essential proteins.

## Results

We assembled the human essentialome constituted by essential genes whose perturbation influences cell viability, ranging from cell death to severe negative impact on cell fitness (*1–11*). We collated 6548 human essential genes/proteins from 13 large-scale experimental studies conducted across various human cell lines [**Table S1**]. Proteins that contain a stretch of five or more identical amino acid runs were classified as proteins with HRs (HRPs), as previously described (*17*). On the basis of essentiality and the presence of HRs, we classified the human proteome into essential proteins with HRs (E-HRPs) and without HRs (E-NonHRPs), and non-essential proteins with HRs (NonE-HRPs) and without HRs (NonE-NonHRPs) [**Figure 1A**]. To understand the influence of essential proteins with HRs, we compared them with E-NonHRPs and NonE-HRPs to assess their influence on phenotypes, molecular interactions, gene expression in space and time and their evolutionary dynamics by assembling and investigating diverse genome-scale datasets [**Table S2**].

**Figure 1.**
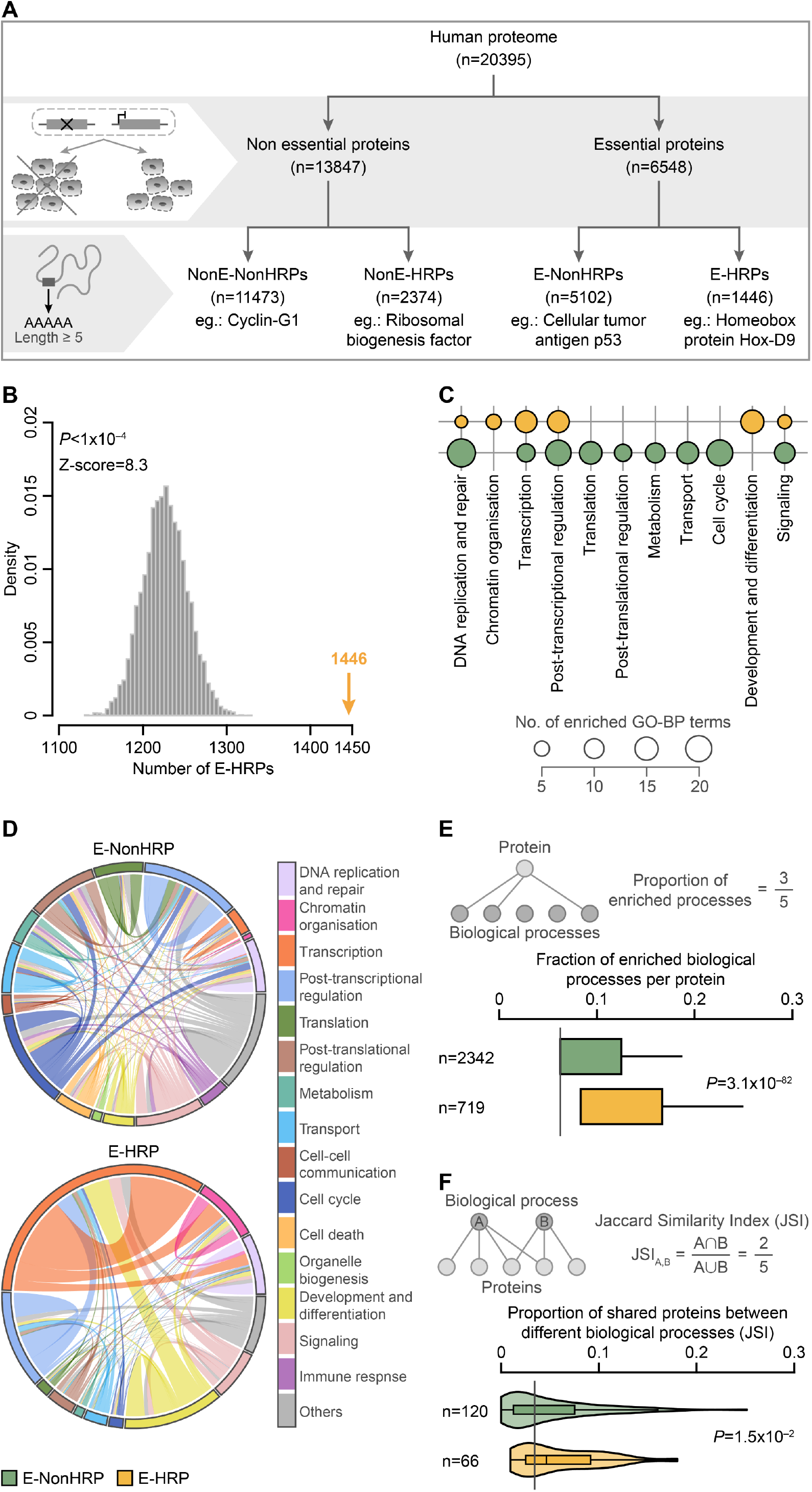
Essential proteins with homorepeats (E-HRPs) are involved in distinct regulatory functions and facilitate cross-talk between processes. (**A**) Classification of the human proteome based on gene-essentiality and presence of amino acid homorepeats. (**B**) Enrichment of HRPs in the human essentialome assessed using permutation testing. The grey histogram represents the random expectation and the yellow arrow the actual observation. (**C**) Bubble plot indicating number of significantly enriched (FDR<0.05) Gene ontology biological process (GO-BP) terms in each of the manually classified major biological processes, with at least three enriched GO terms. (**D**) A functional map highlighting the overlap of E-NonHRPs and E-HRPs across different enriched biological processes. The outer ring represents the biological processes highlighted by different colors, with the arc length corresponding to the number of proteins in each biological process. The thickness of the color is proportional to the number of proteins that are shared between two biological processes. (**E**) Boxplot showing the proportion of enriched biological processes for E-NonHRPs and E-HRPs. Since E-HRPs and E-NonHRPs are involved in a variable number of biological processes, we calculated the proportion of processes each protein is involved in, by normalizing to the total number of processes in that particular class. A higher proportion implies that a protein is involved in more processes. Outliers, beyond the whiskers have not been shown for better visualization. n indicates the total number of proteins in each class. (**F**) Violin plot showing distribution of Jaccard Similarity Index (JSI), which indicates the extent of overlap (top panel) for E-HRPs or E-NonHRPs, between any two biological processes. n represents the number of pairs of biological processes. Statistical significance was estimated using Wilcoxon rank sum test.

### E-HRPs have distinct regulatory functions and bring about cross-talk across biological processes

Essential proteins in humans show an enrichment for proteins with HRs, highlighting the physiological importance of E-HRPs, in higher-order eukaryotes [**Figure 1B**]. E-HRPs show over-representation of polar uncharged (Gln) and charged amino acid HRs (Lys, Asp and Glu), while nonessential HRPs are enriched for non-polar amino acid HRs (Gly, Leu and Pro) [**Figure S1A-S1B**]. Preferential distribution could highlight the importance of the polar amino acid repeats in facilitating key protein functions of E-HRPs. Gene ontology analysis revealed that human E-HRPs are involved in distinct regulatory processes such as chromatin organization, transcriptional and post-transcriptional regulation, and development and differentiation, while E-NonHRPs are preponderantly involved in housekeeping functions such as translation, transport and metabolism [**Figure 1C**]. E-HRPs and E-NonHRPs are involved in shared processes such as transcription, post-transcriptional regulation. Interestingly, within these processes E-HRPs and E-NonHRPs tend to be involved in distinct sub-processes. For instance, (i) in transcription, while E-NonHRPs show enrichment for regulation of transcription mediated by all three RNA polymerases, E-HRPs are predominantly involved in regulation of transcription mediated by RNA polymerase II, (ii) while E-NonHRPs are involved in the post-transcriptional regulation of all types of RNAs, E-HRPs tend to regulate the post-transcriptional processing of mRNAs, specifically alternative splicing [**Figure S1C**]. These findings suggest that E-HRPs predominantly bring about direct regulation of protein coding genes. Although, as a class, E-NonHRPs cumulatively participate in higher number of biological processes compared to E-HRPs (16 and 12 processes, respectively) [**Figure 1D**], at the individual molecule-level, E-HRPs are engaged in higher number of biological processes [**Figure 1E**]. Jaccard Similarity Index (JSI) estimations between different biological processes revealed a higher tendency for E-HRPs to be shared among different processes than E-NonHRPs [**Figure 1F**].

On the other hand, non-essential HRPs enriched for transcription, and development and differentiation show fewer gene ontology terms than those enriched for E-HRPs [**Figure S1D**]. While both E-HRPs and NonE-HRPs participate in the regulation of transcription mediated by RNA polymerase II, E-HRPs are also enriched for regulation of RNA polymerase II mediated transcription of snRNA, suggesting that E-HRPs regulate global splicing events at both transcriptional and post-transcriptional levels. Besides this, within development and differentiation, nonessential HRPs are enriched for anterio-posterior pattern specification, skeletal and nervous system development, while E-HRPs are involved in heart, lung and forebrain development, stem cell population maintenance and *in utero* embryonic development [**Figure S1E**]. Collectively these findings suggest that E-HRPs have distinct functions compared to E-NonHRPs and NonE-HRPs and are multifunctional, affecting cross-talk across various biological processes.

### E-HRPs show high interactability and expansive global regulation

How do E-HRPs affect diverse biological processes? We posit that molecular pleiotropy can be achieved through (i) high interactability and/or (ii) the ability to globally regulate a large number of targets. To examine this, we assembled a comprehensive dataset of experimentally determined molecular interactions spanning functional (protein-protein) and regulatory interactions (protein-DNA and protein-RNA; See **Table S2** for details). E-HRPs show higher number of protein-protein interactions [**Figure 2A; Table S3**] compared to E-NonHRPs. This observation is not confounded by protein disorder content [**Figure S2**]. Analysis of link communities, which represent an array of proteins participating in similar processes, showed that E-HRPs tend to participate in more link communities in the protein interaction network than the E-NonHRPs [**Figure 2B; Table S3**]. Thus E-HRPs act as connectors and bring about cross-talk between different biological processes. Investigation of the protein-complex network, revealed that E-HRPs are constituents of larger multimeric complexes than E-NonHRPs [**Figure 2C**], though both participate in comparable number of protein complexes. Similarly, E-HRPs are members of higher number of phase-separated structures than E-NonHRPs [**Figure 2D**]. Interestingly, E-HRPs are enriched in regulatory condensates such as stress granules, Ribonucleoprotein (RNP) granules, nuclear speckles and paraspeckles, while E-NonHRPs are enriched in postsynaptic density, centrosome and nuclear pore complex, which are primarily involved in housekeeping processes such as cell-cycle, signaling, transport and cytoskeleton organization [**Figure 2E**]. These observations concur with our findings from Gene Ontology enrichment analyses highlighting the regulatory roles and house-keeping functions of E-HRPs and E-NonHRPs, respectively. Network randomization analysis revealed that E-HRPs are enriched for essential protein interactors (both E-HRP and E-NonHRP interactors) and proteins that are inter-community hubs compared to E-NonHRPs [**Figure 2F-2G**]. We did not find any difference in the distribution of intra-community hubs between the two classes of essential proteins [**Figure 2G**]. This implies that the interaction-driven molecular pleiotropy of E-HRPs is not just limited to higher number of interactions but is also determined by the functional importance of their interactors.

**Figure 2.**
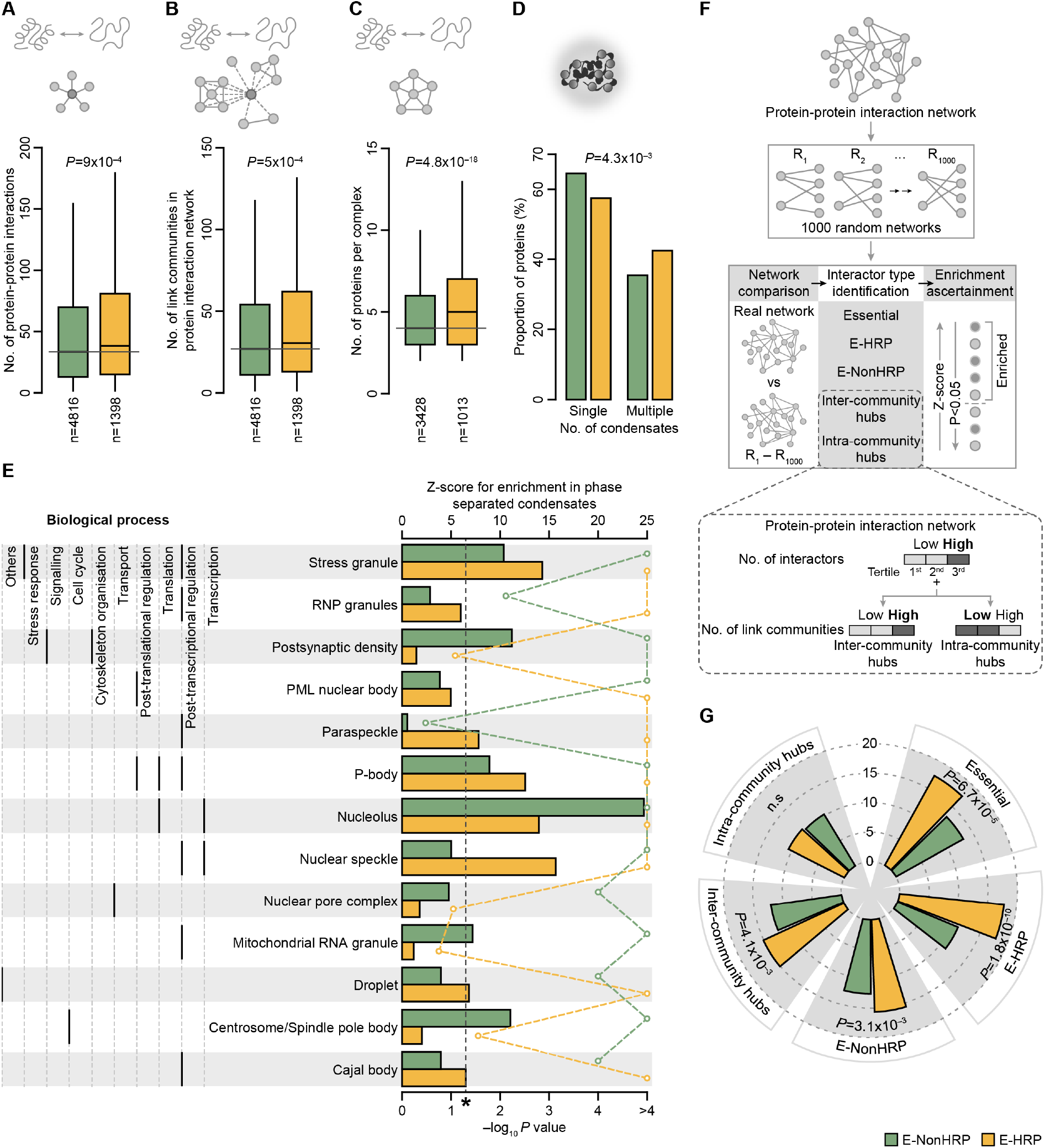
E-HRPs show molecular pleiotropy contributed by their high interactability. Boxplot of distributions of the number of (**A**) protein-protein interactions, (**B**) protein-protein link communities and (**C**) proteins found in a protein complex for E-HRPs and E-NonHRPs. Statistical significance was estimated using Wilcoxon rank sum test. (**D**) Proportion of E-HRP and E-NonHRP proteins found in single or multiple phase separated condensates. n denotes the total number of proteins in each class. P value was computed using Fisher’s Exact test. (**E**) Enrichment of E-HRPs and E-NonHRPs in different phase separated condensates. Condensates with at least 20 proteins were selected for the analysis. Z scores (top axis; bars) and P values (bottom axis; dotted line) were computed using permutation testing. The biological process that each condensate is involved in, was manually mapped, and is shown on the left side of the figure. (**F**) Schematic representation of the methodology adopted for randomization of protein-protein interaction networks. We then checked if the interactor/target of each protein in each random network was an essential protein, E-HRP, E-NonHRP, intercommunity hub or intra-community hub. Z-scores and P-values were estimated by comparing the distribution of the random networks with the observed values from the real network. The inset shows the classification of proteins into inter- and intra-community hubs in protein-protein interaction network. (**G**) Bar plot of distribution of E-HRPs and E-NonHRPs showing enrichment for different types of protein-protein interactors, computed from network randomization. P value was computed using Fisher’s exact test.

We next investigated the regulatory interaction networks to examine the magnitude of genome/transcriptome that E-HRPs regulate. We find that E-HRPs regulate significantly larger number of targets, both at the transcriptional (Transcription Factor-gene target network) and post-transcriptional (RNA-Binding Protein-mRNA target network) levels, compared to E-NonHRPs [**Figure S3A-S3B**]. Importantly, network randomization analysis revealed that E-HRP transcription factors (TFs) are enriched for E-HRP targets, and E-HRP RNA-binding proteins (RBPs) showed a higher tendency to regulate the mRNAs of the essential proteins as well as that of E-HRPs [**Figure S3C-S3E; Table S3**]. These findings imply that E-HRPs not only regulate larger parts of the genome and transcriptome, but also regulate functionally important molecules. Taken together, molecular pleiotropy of E-HRPs can be explained by their ability to (i) engage in more binary interactions, multimeric interactions and higher order assemblies, (ii) regulate relatively larger parts of the genome and transcriptome and (iii) regulate and/or interact with functionally important molecules.

### Diverse factors contribute to the higher interaction potential of E-HRPs

Diverse molecular features such as (i) presence of HRs (*16, 17*), (ii) post-translational modifications (PTMs) (*25, 26*), (iii) eukaryotic linear motifs (ELMs), (iv) alternatively spliced isoforms (*27, 28*), and (v) broad tissue expression profiles (*29*) can influence the protein interaction potential. Bayesian inferences based on log_2_ likelihood ratio (log_2_ LR) estimations revealed that polar (uncharged and charged) and certain non-polar (Ala and Gly) amino acid HR types in E-HRPs showed positive log_2_ LRs for different network attributes that signify higher protein-protein interactability. Compared to nonessential HRPs, almost all amino acid HRs, barring polyCys show positive log_2_ LRs for higher protein-protein interactability for E-HRPs [**Figure 3A**]. For E-HRP TFs and RBPs with higher number of targets, polar HRs showed higher likelihood ratios [**Figure S3F**]. There were no significant differences in the distribution of the structural propensities of the HRs in the E-HRPs and NonE-HRPs, in the limited structural data available in the Protein Data Bank [**Figure S4**]. Taken together, these findings imply that distinct amino acid HRs contribute to higher molecular interactions of E-HRPs.

**Figure 3.**
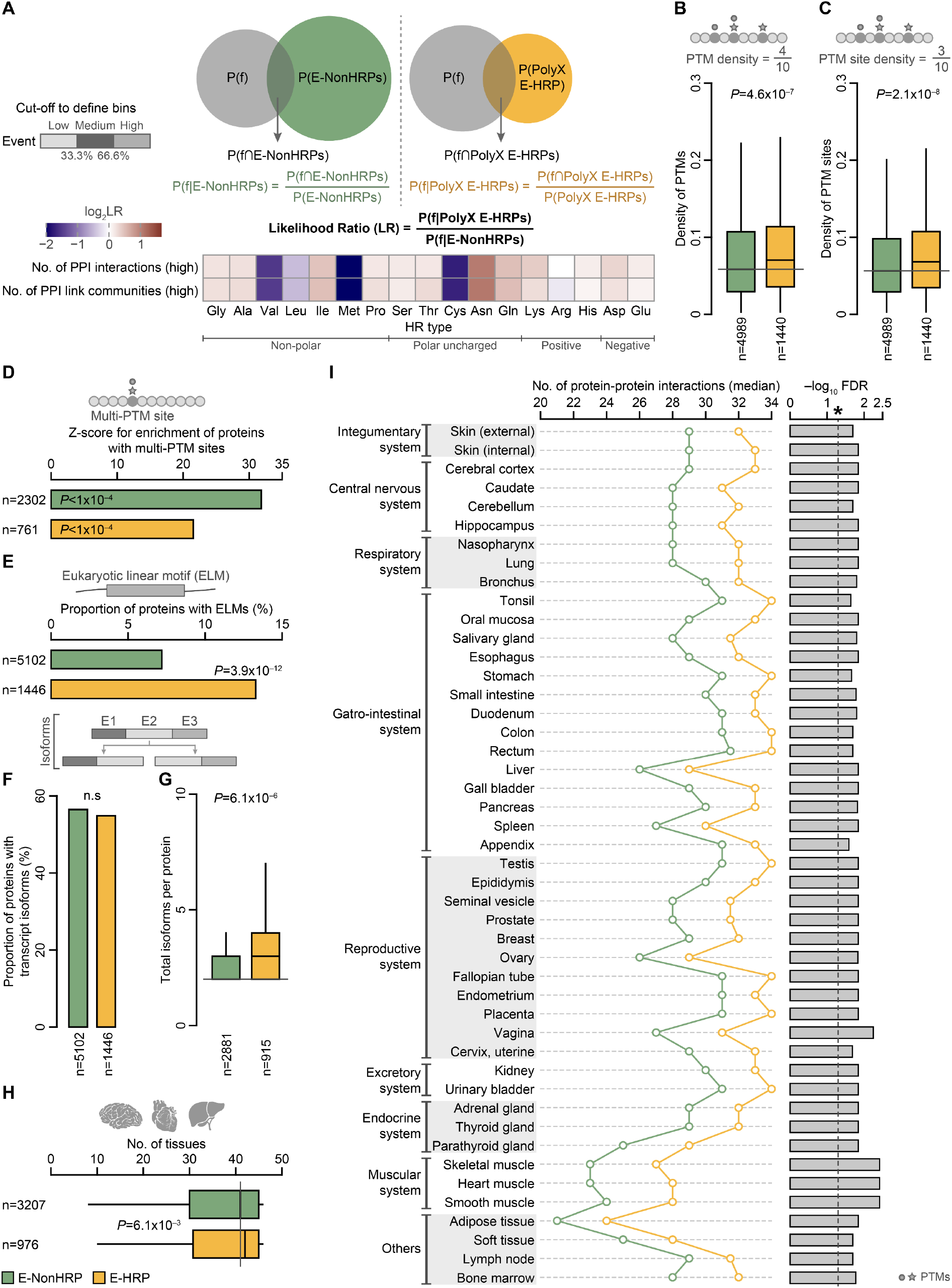
Factors influencing the higher interactability of E-HRPs. (**A**) Likelihood ratios for different repeat types containing E-HRPs for high interactions and link communities in protein-protein interaction network. The top panel presents the schema for Likelihood ratio estimation. Boxplot representing (**B**) PTM density and (**C**) PTM site density for E-HRPs and E-NonHRPs. n denotes the number of proteins in each class. P-value was computed using Wilcoxon rank sum test. (**D**) Enrichment of E-HRPs and E-NonHRPs for protiens with multi-PTM sites. n denotes the number of proteins in each class. Z-scores and P-values were computed using permutation testing. Proportion of E-HRPs and E-NonHRPs (**E**) which harbour Eukaryotic Linear Motifs (ELMs) and (**F**) with alternatively spliced isoforms. n denotes the number of (**E**) proteins and (**F**) proteins with isoforms. P-value was computed using Fisher’s Exact test. (**G**) Distribution of the number of isoforms per protein for E-HRPs and E-NonHRPs. n denotes the total number of proteins which have isoforms. P-value was estimated using Wilcoxon rank sum test. (**H**) Boxplot representing the number of tissue types in which E-HRPs and E-NonHRPs are detected. n denotes the total number of proteins in each class. P-value was estimated using Wilcoxon rank sum test. (**I**) Distribution of the differences in the number of protein-protein interactions of E-HRPs and E-NonHRPs in different tissues. The line plot shows the median distribution of the two classes, and the bar chart shows the significance estimates. P value was computed using Wilcoxon rank sum test and then corrected for multiple testing using FDR.

Importantly, E-HRPs show higher density of PTMs and PTM sites compared to E-NonHRPs [**Figure 3B-3C**]. In addition, the chance of finding PTM sites around the HRs is higher in E-HRPs compared to NonE-HRPs [**Figure S5**]. Based on the number of different types of PTMs found at a PTM site, we classified the PTM sites into those that have (i) only a single modification (mono-PTM sites) and (ii) multiple modifications at the same site (multi-PTM sites). Strikingly, E-NonHRPs are enriched in proteins with multi-PTM sites though they show lower density of PTM sites compared to E-HRPs [**Figure 3D**]. This implies that different regulatory modes could modulate the functionality of different classes of essential proteins. Mono-PTM sites act in consort as rheostat for step-wise functionality (e.g. Cyclin D1; **Figure S6A**). Mono-PTM sites can also allow non-competitive binding of multiple partners to different modifications simultaneously, allowing multivalent binding interactions (e.g. double-stranded repair protein MDC1; **Figure S6B**). This could be one of the factors that aid the participation of E-HRPs in large protein complexes. Contrarily, multi-PTM sites can act as switches for spatio-temporal regulation of protein function and stability, where each of the different PTM states at a given site could encode a distinct function (e.g. Tumor protein p53; **Figure S6C-S6D**). These findings suggest that more PTMs, at different positions, especially around HRs, can aid in multivalent binding or act as rheostats contributing to functional versatility of E-HRPs. On the other hand, limited PTMs in E-NonHRPs, often occurring at the same sites, restrict their binding partner contingent. Interestingly, more E-HRPs contain functional motifs like ELMs (*30*) compared to E-NonHRPs [**Figure 3E**]. These short linear motifs can facilitate the transient and reversible interactions of E-HRPs, bringing about diverse molecular functions.

Although both E-HRPs and E-NonHRPs have similar proportion of proteins with multiple coding isoforms, E-HRPs showed higher number of isoforms per protein compared to E-NonHRPs [**Figure 3F-3G**]. Interestingly, the exons that code for HRs are more often retained in E-HRPs compared to the nonessential HRPs, highlighting that the presence of HRs could be crucial for the functionality of E-HRPs [**Figure S7**]. E-HRPs also show broad tissue-wide distribution compared to E-NonHRPs [**Figure 3H**]. Nevertheless, in each tissue type E-HRPs show higher number of interactors compared to E-NonHRPs [**Figure 3I**]. Collectively these findings suggest that the increased proteome diversity and extensive retention of HRs, higher PTM density and broad tissue expression contribute to higher interaction potential of E-HRPs.

### E-HRPs influence human-specific embryonic and brain development

One of the fundamental biological processes that human E-HRPs are enriched in, is development and differentiation [**Figure 1C**]. What is the influence of E-HRPs on human development? Dynamic changes in the temporal transcriptomic profiles of genes, which promotes rapid rewiring of molecular networks, are a key hallmark of development. Such dynamic alterations facilitate cell-fate specification and cell lineage determination. To understand the role of E-HRPs in human development, we first examined genes that are developmentally dynamic (DDGs) (*31*) across different organs, derived from all the three germ layers (ectoderm, mesoderm and endoderm). Developmental dynamicity is a phenomenon wherein genes exhibit extensive alterations in the expression levels across different stages of development (*31*). E-HRPs are developmentally dynamic across a higher number of organs compared to E-NonHRPs [**Figure 4A**]. Notably, across organs, E-HRPs show lower tissue specificity, implying that they are expressed in more tissues, compared to E-NonHRPs [**Figure 4B**]. These findings indicate that E-HRPs are expressed dynamically across a broad spectrum of tissues and organs. This highlights the importance of the dynamic tuning of E-HRP expression and its downstream consequences in human development. Strikingly, E-HRPs are more developmentally dynamic in the ectoderm-derived human forebrain and hindbrain [**Figure 4C**]. Such developmental dynamicity of E-HRPs in the forebrain and hindbrain is evolutionarily conserved across mammals [**Figure S8**]. E-HRPs are flanked more by human accelerated regions (HARs) [**Figure 4D-4E**] and such HAR-flanking is more pronounced for E-HRP TFs [**Figure 4F**]. HARs are evolutionarily conserved sequences in mammals but are fast evolving in humans, and these sequences overlap with regulatory protein binding motifs such as transcription factor binding sites (*32*). This implies that the E-HRPs, which are predominantly gene-expression regulators, are regulated in a species-specific manner in humans. While HAR-flanked E-HRPs and E-HRP TFs show enrichment for embryonic and brain development [**Figure S9A-S9B**], HAR-flanked E-NonHRP TFs were enriched for stem cell differentiation, and forelimb development [**Figure S9B**]. E-HRPs [**Figure 4G; top panel**] along with forebrain and hindbrain E-HRP DDGs [**Figure 4G; bottom panel**] are predominantly flanked by human acceleratory regions (HARs) active in brain development. This suggests that the expression levels of E-HRPs that are involved in development are regulated in a human-specific manner.

**Figure 4.**
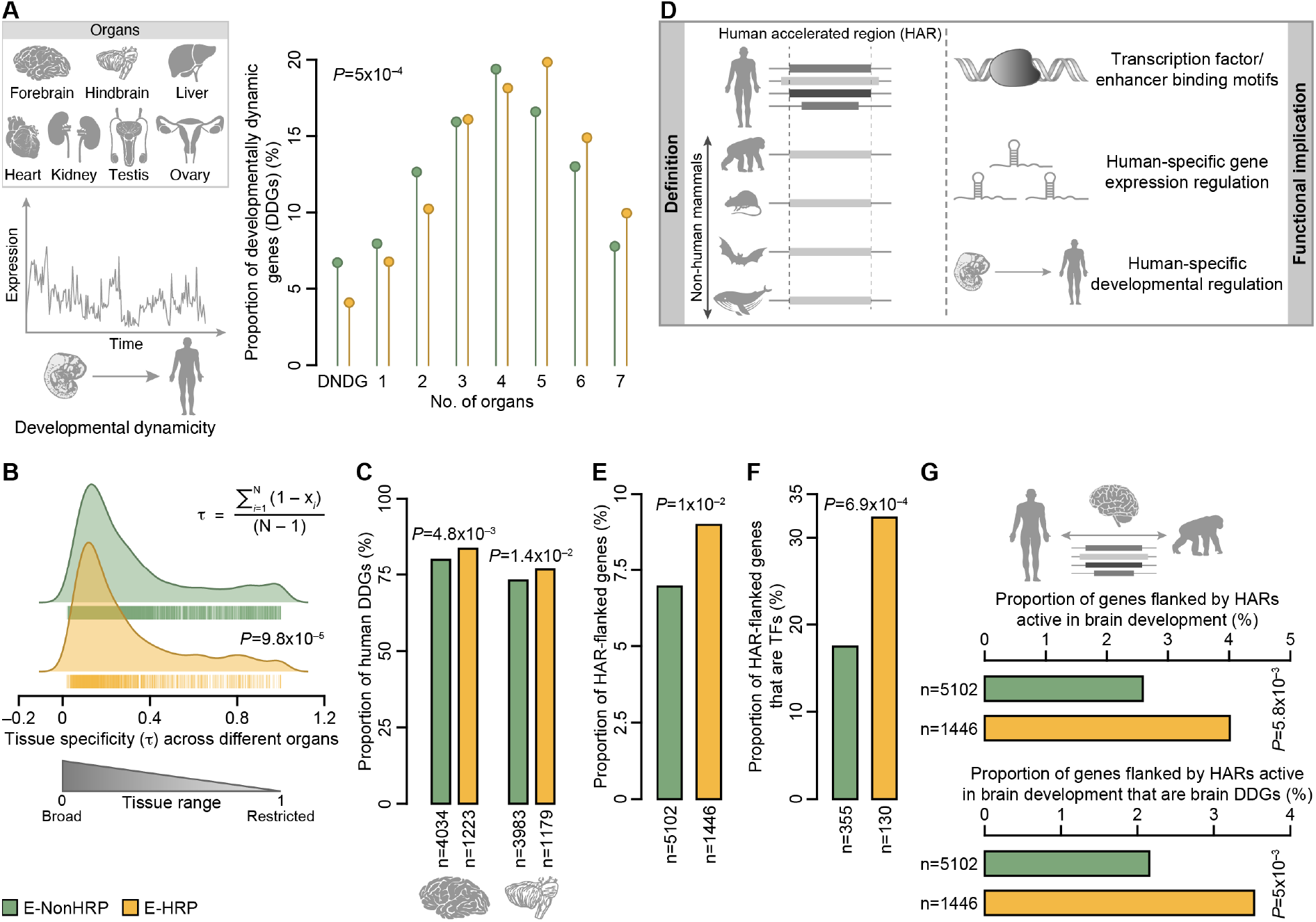
E-HRPs are developmentally dynamic across multiple organs across different species. (**A**) Proportion of developmentally dynamic genes (DDGs) among E-HRPs and E-NonHRPs in different number of organs. The organs in which DDGs were identified are presented in the top left panel. P-value was computed using Fisher’s Exact test. DNDG represents developmentally non-dynamic genes. (**B**) Tissue specificity of E-NonHRPs and E-HRPs. The area of the curves represents the density of the distribution. The bars at the bottom of the density distribution represent the data points. P-value was computed using Wilcoxon-rank-sum test. N is the total number of organs and x_i_ is the maximum expression in each organ in the course of development normalised with the maximum expression across organs (*31, 74*). (**C**) Distribution of E-HRP and E-NonHRP DDGs in the ectoderm-derived forebrain and hindbrain. n denotes the number of genes in each class in each organ. P-value was estimated using Fisher’s Exact test. (**D**) Illustration showing definition and functional implications of human accelerated regions (HARs). Proportion of (**E**) genes flanked by HARs, (**F**) HAR-flanked E-HRP and E-NonHRP TFs. n denotes the number of genes in each class. P-value was computed using Fisher’s Exact test. (**G**) Proportion of genes flanked by HARs that are active in brain development (top panel) and those that are active in brain development that are DDGs in human forebrain and hindbrain development (bottom panel). n denotes the number of genes in each class. P-value was computed using Fisher’s Exact test.

Since E-HRPs are enriched for regulatory processes, we next aimed to delineate the temporal regulatory roles of E-HRPs and their influence on human development. To address this, we investigated the systems-level effects of E-HRPs in different stages of human pre-gastrulation and prenatal brain development [**Figure 5A**]. For this, we collated developmental genes from publicly available single-cell RNA sequencing data of human prenatal developmental stages [**Table S2**]. E-HRPs regulate higher number of gene targets compared to E-NonHRPs in the pre-gastrulation stages [**Figure 5B**]. Interestingly, we find increased gene regulation by E-HRPs in specific brain development stages such as gestation week (GW) 8, 9, 13 and 19 [**Figure 5C**]. Strikingly, these time points correspond to cell differentiation and cell type specification into (i) frontal, parietal, temporal and occipital lobes (GW8 and GW9), (ii) cell migration of newly formed neurons in the cortex (GW13), and (iii) axonogenesis and formation of connections between neurons (GW19) of brain (*33, 34*). This could explain the importance of E-HRPs in regulating the key developmental transitions. Interestingly, at the post-transcriptional level also, E-HRPs tend to regulate more number of transcripts compared to E-NonHRPs in both human pre-gastrulation and brain development stages [**Figure 5D-5E**]. Exemplifying the aforementioned trends are the developmentally important HOX, FOX and SOX cluster of genes. These clusters contain E-HRPs, some of which tend to be developmentally dynamic and are flanked by HARs [**Figure 5F**]. Furthermore, E-HRPs show a higher number of protein-protein interactions across different pre-gastrulation and brain developmental stages [**Figure S10**]. Taken together, human-specific regulation of E-HRPs, which physically interact and/or regulate larger portions of developmental genome and transcriptome, might drive processes underlying human-specific embryonic and brain development.

**Figure 5.**
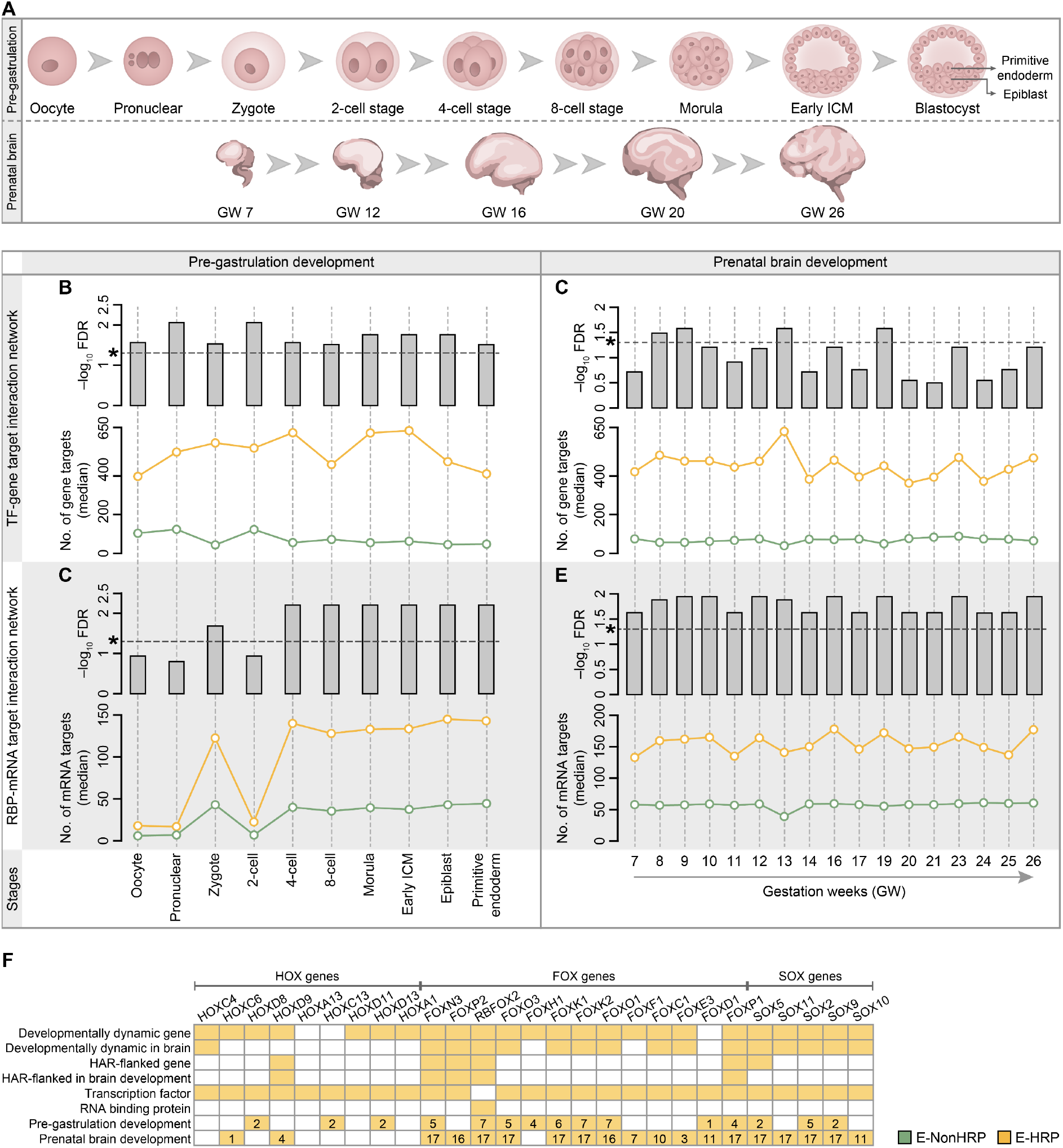
E-HRPs are involved in modulating temporal regulation of prenatal development. (**A**) Illustration of human pre-gastrulation and prenatal brain developmental stages studied here. Distribution showing median number of interactors/targets of E-HRPs and E-NonHRPs during different stages of pre-gastrulation and prenatal brain development (bottom panel) and its corresponding significance estimates (top panel) for TF-target regulatory network (**B** and **D**) and RBP-RNA regulatory network (**C** and **E**), respectively. The different developmental stages are given in the X-axes. Correction for multiple testing to estimate significance was done using FDR. The significance line is denoted using an asterisk, with the values above the asterisk representing statistical significance. ICM stands for Inner cell mass. (**F**) Attributes of important regulatory E-HRPs involved in pre-gastrulation and pre-natal brain development, exemplifying human-specific regulation of development. The numbers in each cell provide the total number of stages, investigated here, in which the particular gene is expressed.

### E-HRPs constitute the rapidly evolvable part of the essentialome

Given the human-specific functional importance of E-HRPs, we next aimed to examine if E-HRPs have different evolutionary trajectories compared to E-NonHRPs. Estimation and analysis of the evolutionary age of human proteins across 460 eukaryotes [**Figure S11**] revealed that E-HRPs tend to show a clade-specific expansion in chordates and metazoans, whereas, E-NonHRPs are predominantly pan-eukaryotic [**Figure 6A**]. When we investigated the biological processes these proteins are involved in, we observed that the clade-specific expansion of E-HRPs corresponds to an enrichment in transcription and development and differentiation related processes [**Figure 6B**]. This implies that E-HRPs might have originated and co-evolved with the emergence of (i) organismal complexity (e.g., multicellularity) in metazoans and (ii) notochord and/or brain in chordates.

**Figure 6.**
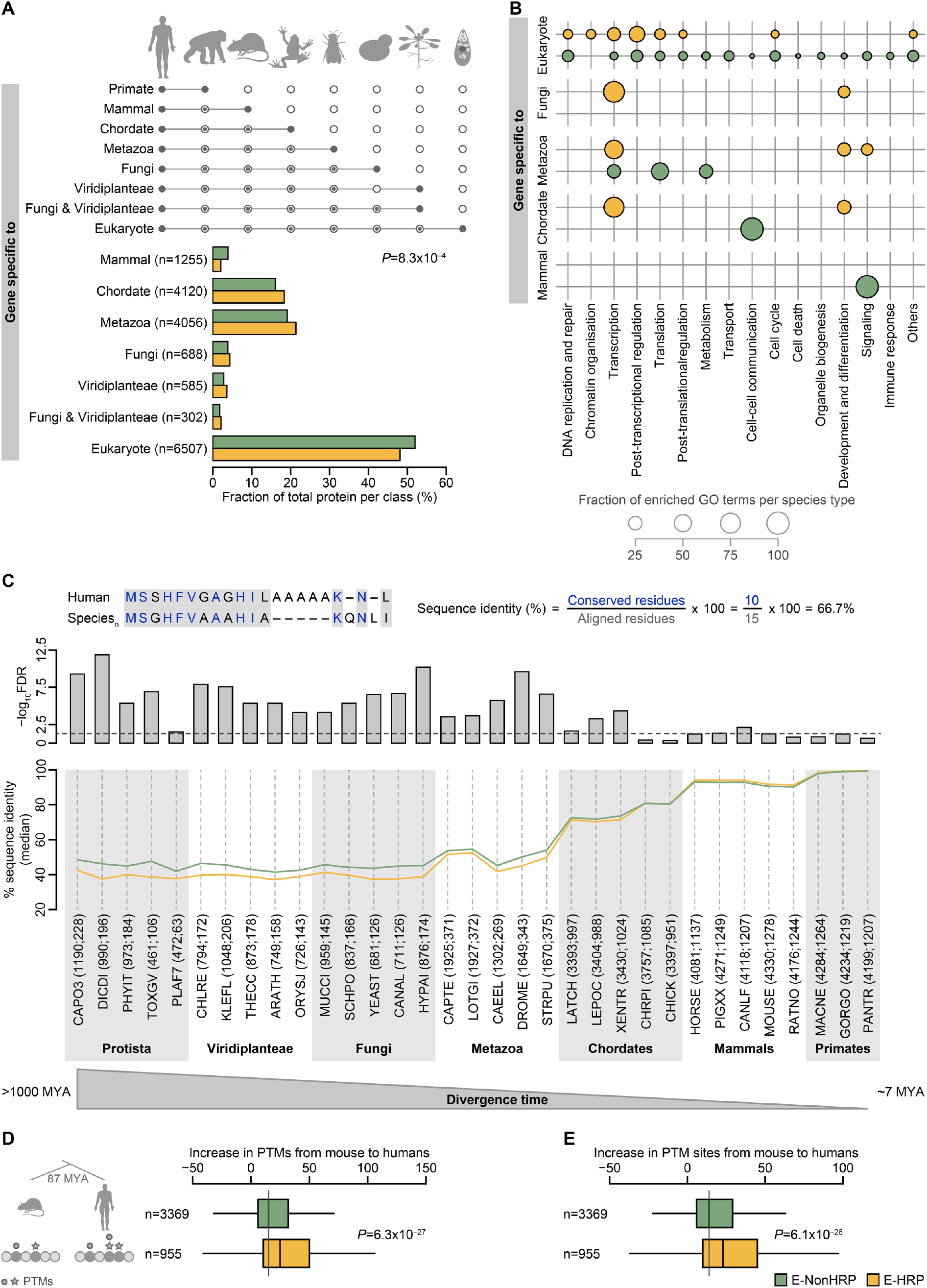
E-HRPs show lineage-specific expansion and evolve rapidly. (**A**) Estimation and distribution of the evolutionary age of E-HRPs and E-NonHRPs. n denotes the total number of proteins in each taxon. P-value computed using Chi-squared test. (**B**) Bubble plot indicating number of significantly enriched (FDR<0.05) Gene Ontology biological process (GO-BP) terms in each of the manually classified broad biological processes. The size of the bubble denotes the percentage of enriched GO-BP terms in each category. (**C**) Median of percentage sequence identity (bottom plot) of human sequences and their orthologs across 33 species (X-axis) and their significance estimates (top plot). P-value was computed using Wilcoxon-rank-sum test and multiple hypothesis correction was done using FDR. The taxonomic details of the abbreviated species provided in the X-axis are given in **Table S4**. Boxplot showing increase in (**D**) PTMs and (**E**) PTM sites in E-HRP and E-NonHRP proteins from mouse to humans. n denotes the number of 1:1 protein orthologs in each class. P-value was computed using Wilcoxon-rank-sum test.

We next aimed to examine the extent of sequence divergence of the two classes of essential proteins. For this, we computed the percentage sequence identity of every human protein, with their respective one-to-one orthologs across 33 representative species spanning different taxonomical classes (∼1000 Mya of evolutionary divergence time). Strikingly, E-HRPs showed about 3.2% lesser sequence identity across primates to protists, compared to E-NonHRPs [**Figure 6C**]. This means that for the average protein length of 500 amino acids, E-HRPs would accumulate about 16 amino acid substitutions more than E-NonHRPs of the same length. Such increased sequence divergence reflects that essential proteins with HRs rapidly evolve compared to those without HRs. To assess the functional implications of such elevated sequence variation in E-HRPs, we investigated the one-to-one orthologs of human proteins in mouse (∼87 million years divergence time) and computed the difference in PTM sites in human essential proteins compared to their mouse orthologs. Interestingly, the human E-HRPs showed an increased acquisition of PTMs and PTM sites than the E-NonHRPs [**Figure 6D-E**]. Furthermore, there is a higher tendency to find mutations in the PTM sites in the E-HRPs leading to diseases than E-NonHRPs, highlighting their functional importance [**Figure S12**]. Collectively, these findings suggest that the rapid divergence of E-HRPs might aid in accelerated and increased accumulation of functionally important sites in E-HRPs, thereby contributing to the expansion of the functional spectrum of E-HRPs.

## Discussion

Systems-level studies so far have focused on elucidating the functional attributes of essential proteins compared to non-essential proteins (*1–11, 35–50*). Here we resolve the paradox of the prevalence of the hypermutable HRs in essential proteins, which are known to have conserved functionalities, by delineating the functional and evolutionary attributes of essential proteins with HRs. The findings presented here highlight that essential proteins with HRs tend to be highly interactive, bring about cross-talk between diverse processes and are involved in distinct regulatory functions, including that of human-specific processes such as development [**Figure 7**]. Factors such as more PTMs and presence of HRs contribute to such functional versatility. Molecular studies show that HRs in essential proteins have varying functional roles. For instance, (i) the polySer HR in the human essential protein JMJD6 aids in localization to the nucleus (*20*), which is vital for this enzyme with dual functions of arginine demethylase and lysl-hydroxylase to bring about the regulation of transcription and pre-mRNA splicing and (ii) polyHis HR in human DRYK1A aids in its localization to nuclear speckles which is essential for its role in RNA synthesis and processing (*21*). We note that although the systems-level analyses provide general trends, supported by specific molecular instances, every general trend reported here might not be applicable to all E-HRPs.

**Figure 7.**
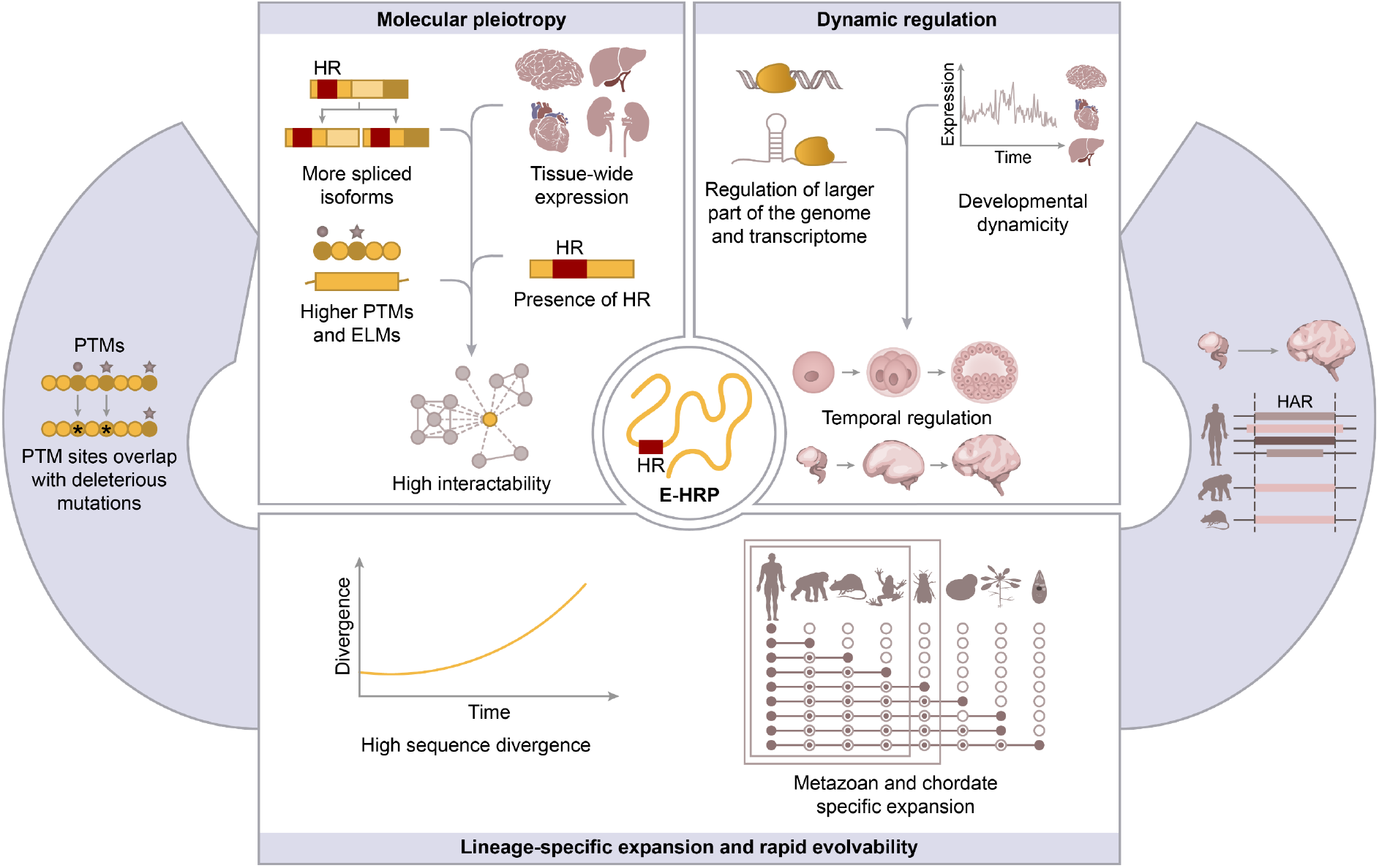
Impact of the essential proteins with homorepeats on fitness. E-HRPs (i) show molecular pleiotropy (top left), (ii) affect dynamic temporal regulation of human development (top right), (iii) tend to show rapid sequence divergence, amounting to gain of functionally relevant PTMs (bottom left) and (iv) show clade-specific expansion and with human-specific regulatory elements dynamically regulate human development (bottom right).

In addition to the functional contributions, the presence of HRs also provides evolutionary benefits to the essential proteins. Our data highlights that HR-associated genetic variation can lead to gain of functionality through the acquisition of functional sites such as PTMs [**Figure 7**]. Additionally, synonymous genetic variation in proteins might also lead to expression effects (*51*). Since E-HRPs act as hubs in regulatory networks, modulating their activity or levels through HR length variation or single nucleotide changes can result in accelerated evolution through network rewiring, contributing to adaptability (*18, 52*). This implies that essential proteins with HRs constitute rapidly evolvable essentialome, a systems-level attribute that is conserved across yeast and human (*17*). Budding yeast HRPs are more prevalent in gene duplicates (*17*) and gene duplication is often perceived as a contributor to relaxed selection (*53*). However, in humans, only one-fourth of the E-HRPs tend to contain gene-duplicates [**Figure S13**], thereby highlighting that the rapid sequence divergence of E-HRPs could be associated with presence of HRs rather than genetic compensability/redundancy. These findings imply that the presence of HRs in essential proteins might contribute to the trade-off between robustness and evolvability.

Our data demonstrates that, besides contributing to evolvable essentialome, proteins with HRs contribute to the emergence of human-specific processes such as embryonic and brain development. Remarkably, other human-specific elements such as HARs tend to flank E-HRPs, bringing about human-specific regulation. This concerted action between different genomic, transcriptomic and proteomic features could cause a butterfly effect, impacting diverse human-specific physiological outcomes. Emergence of low-complexity regions in proteins is a prominent evolutionary mechanism found in novel proteins and fast-evolving, young protein families (*54, 55*). Our results corroborate with these findings and further extend this paradigm by showing that essential proteins with LCRs, represented in its purest form by homorepeats, could regulate species-specific processes. Is the enrichment of proteins with HRs in essentialomes an evolutionarily conserved phenomenon? While the essentialomes of prokaryotes and fission yeast are not enriched for E-HRPs, those of Arabidopsis, Zebrafish and mouse alongside budding yeast and humans are enriched for proteins with HRs [**Figure S14**]. This suggests that the enrichment of E-HRPs in essentialomes could be an after-effect of whole genome duplication events (*56–60*), which correspond to emergence of complexity and diversity. The systems-level conservation of enrichment of E-HRPs in essentialomes signify that our findings in humans presented here could provide insights for understanding the role of essential proteins with HRs in other species as well.

Not surprisingly, abnormal repeat expansions in such functionally important E-HRPs can lead to diseases. For instance, polyGln expansion in Huntington protein (HTT) causes conformational changes in the protein and leads to neurodegenerative Huntington disease (*61*) and polyAla HR in human Runt-related transcription factor 2 (RUNX2), a transcription factor involved in osteoblast differentiation and skeletal morphogenesis, can lead to protein aggregation and impair protein functionality, leading to cleidocranial dysplasia in humans (*62*). The results presented here highlight the potential of targeting the E-HRPs for therapeutic intervention. For instance, in neurological diseases (i) inhibition of the E-HRP, C-Abelson (c-Abl) tyrosine kinase confers neuroprotection by abrogating dopaminergic neuronal death, thereby alleviating Parkinson’s disease (*63*), (ii) targeting PTM sites acquired in pathological conditions, as shown in the case of Huntington’s disease, could be a potential intervention strategy (*64*). Furthermore, in cancers, the E-HRP (i) Epidermal Growth Factor Receptor (EGFR) targeting by monoclonal antibodies and small molecule tyrosine kinase inhibitors has implications in treating colorectal, head and neck cancer and pancreatic cancer (*65*) and (ii) Serine-threonine kinase mammalian target of rapamycin (mTOR) is a validated therapeutic target for renal cell carcinoma (*66*). Therefore, the involvement of E-HRPs in regulation of development and species-specific processes presents opportunities for identifying potential therapeutic targets against different pathologies such as cancers, neurological disorders and infections.

## Materials and methods

### Assembly of human essentialome and identification of amino acid homorepeats

Human essential proteins were identified from experimentally validated studies done across different cell types and in different culture conditions (*1–11*). We identified all genes reported to affect cell survivability in given culture conditions as essential genes. These genes were then mapped to their Uniprot SwissProt accession IDs and finally collated to represent the human essentialome [**Table S1; Supplementary data table 1**]. All other human genes were considered as non-essential. The sequences of all human proteins were obtained from the UniProt SwissProt database. An in-house Perl script was used to identify homorepeats (HR) in the human proteome. A protein was considered as an HRP if it had atleast one HR stretch with a minimum length of 5 residues, as previously defined (*17*). The compendium of the different datasets analyzed in this study, with details on the size of the datasets and thresholds used, if any, are provided in **Table S2**.

### Gene ontology enrichment analysis

Enrichment of Gene ontology biological processes (GO-BP) was estimated using DAVID server using an FDR cut-off of < 0.05 (*67, 68*). The enriched GO-BP terms were then manually organized into major biological processes [**Supplementary data table 1**]. We computed the extent of overlapping genes between any two enriched processes using Jaccard Similarity Index, which represents the ratio of the number of genes common to both biological processes to the sum of all genes in the two processes.

### Network Analysis

We computed the topological properties of protein-protein interaction, Transcription factor-gene target and RNA binding proteins-mRNA target networks such as degree and outdegree using Cytoscape and/or R. We identified the link communities in protein-protein interaction network that each protein participated in using a previously reported algorithm (*69*). Network randomization for protein-protein interaction network was done by generating 1000 random networks using an in-house written Python script. In each random network, the total number of interactions of each protein (degree) was kept constant and the interacting partners (edges) were randomized. For TF-gene target and RBP-mRNA target networks, 1000 random networks were generated using BiRewire package in R (*70*). In every random network, the total number of targets of TFs/RBPs (outdegree) and the number of regulating proteins for gene targets/mRNA (indegree) were kept constant, but the interactions (edges) were randomized.

### Quantification of homorepeat retention in isoforms

Human protein isoform sequences were retrieved from Uniprot, and an in-house Perl script was used to identify homorepeats (HRs) in them. For every HRP, the isoforms were classified as HR-isoform(s) and NonHR-isoform(s). HR retention fraction is defined as the ratio of the number of HR-isoforms to the total number of isoforms of a protein, including the canonical isoform.

### Reconstruction of spatio-temporal networks

We reconstructed tissue-specific and developmental stage-specific networks for protein-protein and/or TF-gene target and RBP-mRNA networks. Tissue-wide expression of proteins pertaining to 46 different human tissues were obtained from Human Protein Atlas (*71*). We reconstructed tissue-specific networks by integrating the global protein-protein interaction network with the tissue-specific proteome information. Human pre-gastrulation and prenatal brain developmental proteomes were collated from high-throughput studies [**Table S2**]. For every stage, stage-specific proteomes were overlaid on the global human protein-protein/ TF-target/ RBP-RNA interaction networks to arrive at the stage-specific interactomes.

### Estimation of PTM site gain in humans

To assess for the PTM site gain in human essential proteins, we retrieved the human and mouse PTMs from dbPTM [**Table S2**]. We then mapped these PTMs on human proteins with 1:1 mouse ortholog (obtained from OMA browser). Increase in the number of PTM sites in human orthologs compared to that of mouse was computed for every pair of orthologs.

### Estimation of evolutionary age of human proteins

We obtained the orthologs of human proteins across 460 eukaryotes [**Figure S10**] from OMA browser (*72*) and retained only 1:1 orthologs across other species. For species with multiple strains, we ensured that the strain with the maximum number of proteins or the most commonly studied model strain was selected as the representative.

We classified the species into distinct taxonomic lineages based on classification obtained from NCBI taxonomy browser (*73*) and manual curation. Based on this information, the species were classified into different taxonomical classes, namely, primates (all primates except human), mammals (all mammals excluding primates), chordates (all chordates excluding mammals), metazoans (all metazoans excluding chordates), fungi, viridiplantae and protists. Every human protein with one-to-one ortholog was classified to be specific to any of the above classes, based on the class in which the most ancestral ortholog could be detected. The human proteins for which the orthologs were detected in both fungi and plants were classified as fungi-viridiplantae specific.

### Estimation of sequence divergence

We considered 33 representative species spanning across different clades from protists to primates for this analysis. Every human protein was aligned in a pairwise manner to the corresponding one-to-one ortholog across the species. Percentage sequence identity (PID) was computed as the ratio of number of conserved residues over the number of aligned residues for every pairwise alignment (PID2) in Biostrings package in R. This ensured that the sequence divergence is not over-estimated due to the absence of a HR in any orthologs. We compared the distribution of the PID of E-HRPs orthologs with that of E-NonHRP orthologs for each representative species and assessed the statistical significance using Wilcoxon rank sum test. The P-values were corrected for multiple testing using FDR.

### Statistical analysis

Statistical significance in the differences in the distribution of discrete variables was assessed using Chi-squared or Fisher’s exact test and that of continuous variables was estimated using the non-parametric Wilcoxon rank sum test. Correction for multiple testing was done using the with the Benjamini–Hochberg (FDR) method for comparisons between the classes for datasets that represent similar biological features. Enrichment of proteins in different classes (for instance, HRPs in essential proteins, E-HRPs in various phase separated structures) was examined using permutation tests by performing 10,000 randomizations. In each permutation, each protein of interest (e.g., E-HRP) was replaced with a random gene and the number of random genes that overlapped with a specific class of proteins (e.g., proteins that participate in a specific phase-separated condensate) was noted for all the randomizations. Using the distribution from the random expectation, Z-score was estimated. Z-score represents the magnitude of deviation of the actual observation compared to the mean of random distribution in terms of the number of standard deviations. We estimated P-values for enrichment as the ratios of the number of the randomly observed proteins ≥ to the number of actually observed proteins to the total number of randomized samples (10,000). For obtaining Bayesian inferences, we classified proteins into those that had low, medium or high molecular interactions, based on tertile cutoffs. We then estimated conditional probabilities for finding high interactability in a molecular interaction network given that the essential protein contained a homorepeat, and for finding a protein with a homorepeat given that it is high interactive. To assess the extent of contribution of different amino acid HR types for high interactability of E-HRPs, we computed likelihood ratio (LR) estimates based on the conditional probabilities, as the ratio of the probability of finding an essential protein with high interactability, given that it contained a specific amino acid HR type divided by the probability of finding the specific feature given that the essential protein did not contain an HR. All statistical analyses were performed in R.

## Acknowledgements

We thank P. L. Chavali, A. Dhayalan and A. D. Allu for comments on the manuscript and R. V. Kadumuri for helping with protein disorder predictions. Funding from IISER Tirupati (to A.K.S., K.S.K., S.C.), Prime Minister’s Research Fellowship (to A.K.S.), Ramalingaswami Re-entry Fellowship (BT/RLF/Re-entry/05/2018 to S.C., I.A.), Junior Research Fellowship (to K.S.K.) from Department of Biotechnology, Start-up Research Grant (SRG/2019/001785; to S.C.) from Science and Engineering Research Board and Kishore Vaigyanik Protsahan Yojana Fellowship from Department of Science and Technology (H.R.), Government of India is gratefully acknowledged.

## Author Contributions

A.K.S. was involved in data collection, data analysis, interpretation, preparation of the figures and writing the manuscript; I.A. was involved in data collection of tissue-specific proteomes, human pre-gastrulation and prenatal brain developmental stage proteomes and analysis of the molecular networks, and analyzing essentialomes of other species; H.R. was involved in generating 1000 randomised networks of protein-protein interactions, computing gene age and percentage sequence identities of essential proteins; K.S.K. was involved in assembling human and mouse PTM dataset, assembling and analysis of PDB structures of human HRs; S.C. was involved in designing and supervising the study, interpreting the data and writing the manuscript. All authors were involved in editing the manuscript.

