## Supplementary Information for "Proteins with amino acid repeats constitute rapidly evolvable and human-specific essentialome"

*Correspondence to:

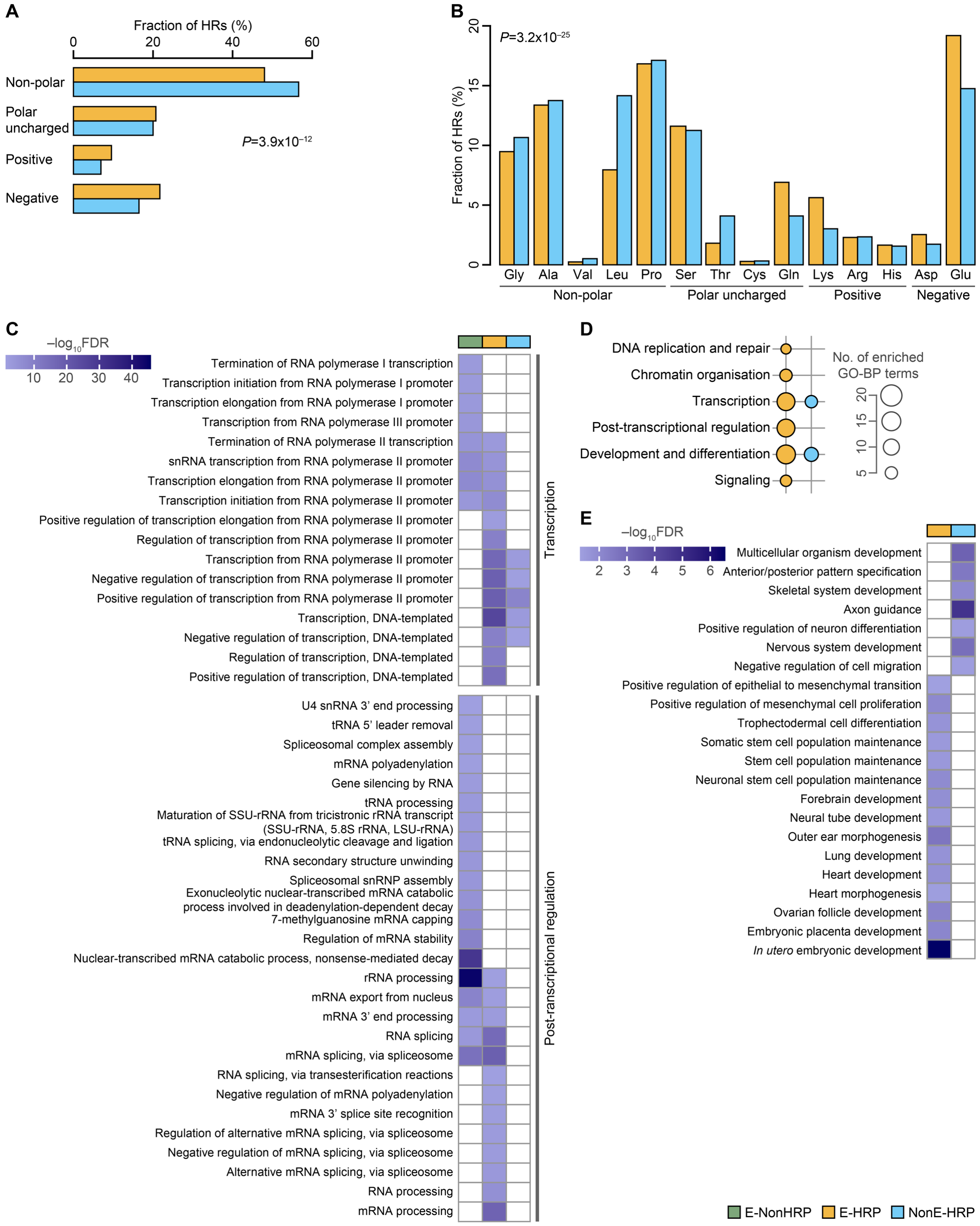


**Fig. S1. Amino acid HR type and functions of essential and non-essential HRPs.** Fraction of amino acid HR types based on (**A**) physicochemical properties and (**B**) different amino acids among E-HRPs and E-NonHRPs. Statistical significance was estimated using Chi-squared test. We considered amino acid HRs with ≥ 5 HR instances in both E-HRPs and NonE-HRPs for this analysis. (**C**) Heatmap showing significantly enriched (FDR<0.05) Gene ontology biological process (GO-BP) terms categorized under transcription and post-transcriptional regulation for E-NonHRPs, E-HRPs and NonE-HRPs. (**D**) Bubble plot representing the number of significantly enriched (FDR<0.05) GO-BP terms in each of the manually classified broad biological processes for E-HRPs and NonE-HRPs. The bubble size denotes the number of enriched GO-BP terms in each category. Only broad processes consisting of at least three enriched GO terms are shown. (**E**) Heatmap showing significantly enriched (FDR<0.05) Gene ontology biological process (GO-BP) terms categorized under development and differentiation for E-HRPs and NonE-HRPs.

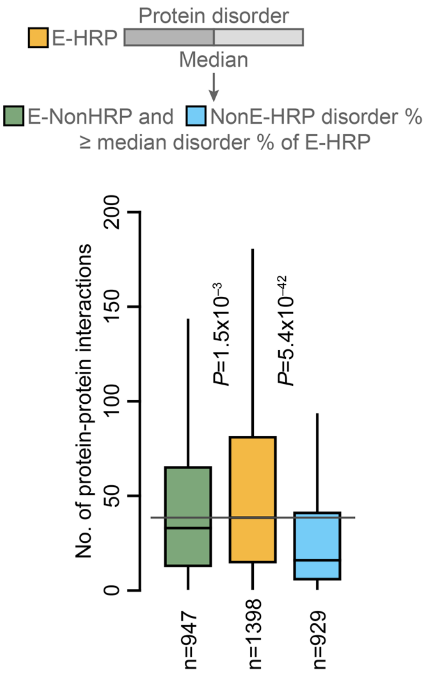


**Fig. S2. Protein disorder does not confound our observations on the high interactability of E-HRPs.** n is the number of proteins of E-HRPs, and the number of E-NonHRP and NonE-HRP proteins with disorder percentage ≥ median of E-HRPs. Statistical significance was estimated using Wilcoxon rank sum test.

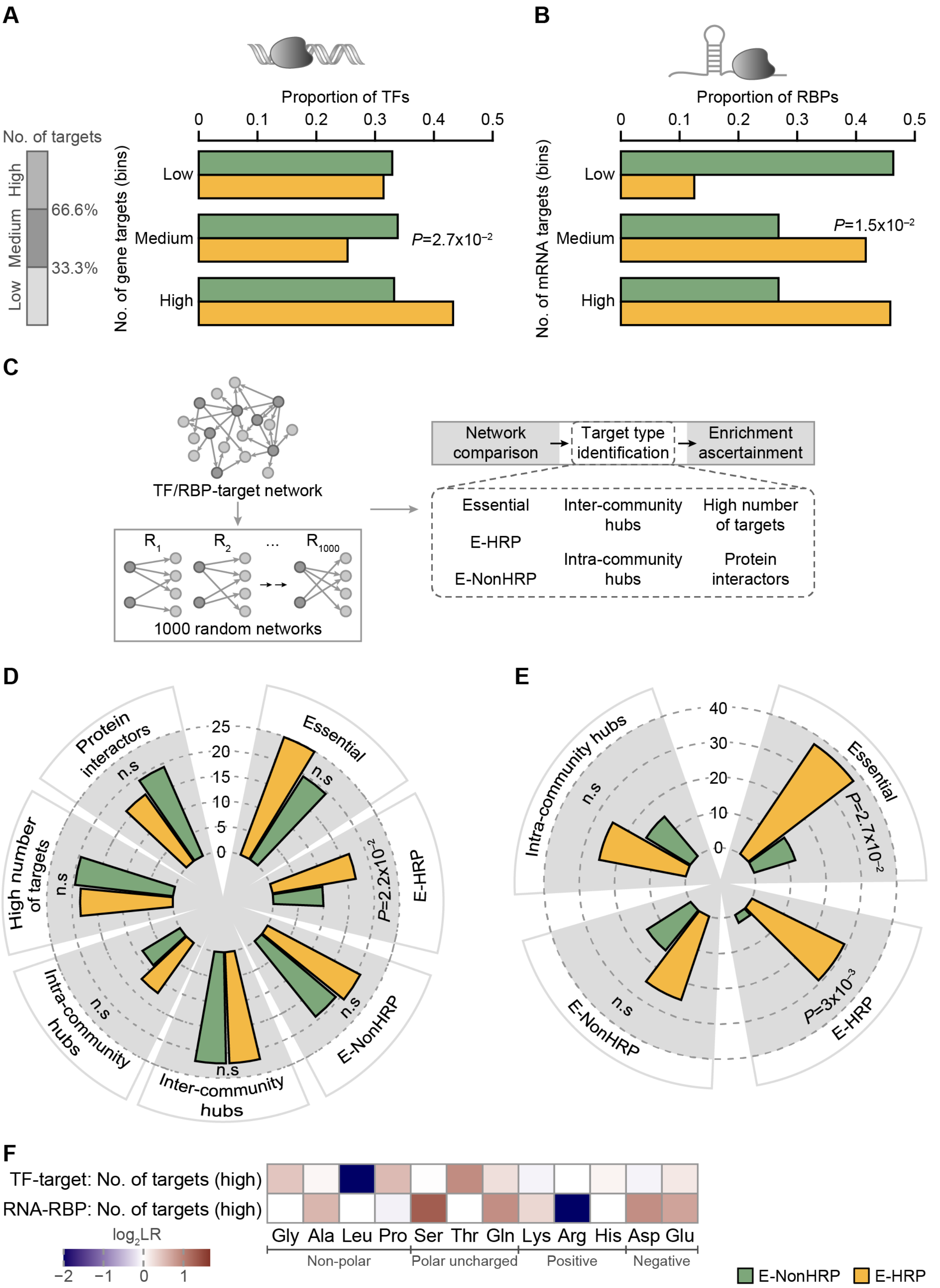


**Fig. S3. E-HRPs regulate larger parts of the genome and transcriptome.** Bar plot of distributions of the number of targets regulated by E-HRP and E-NonHRP (**A**) TFs and (**B**)

RBPs. (**C**) Schema showing generation of random networks for directional graphs and computation of Z-score and P-values using permutation testing. Distribution of E-HRP and E-NonHRP regulators showing enrichment for different types of targets of (**D**) TFs and (**E**) RBPs, estimated from network randomization. P-value was computed using Fisher's exact test. (**F**) Likelihood ratios for different amino acid HR types (X-axis) containing E-HRPs for high gene and mRNA targets of TFs and RBPs, respectively.

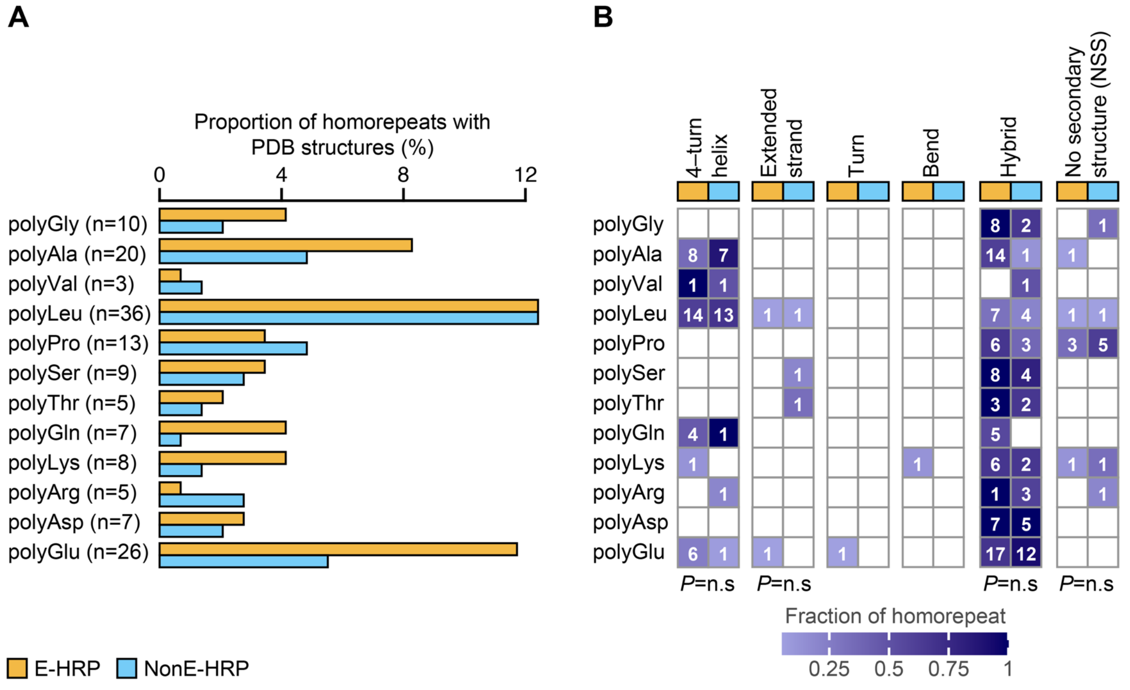


**Fig. S4. Structural propensities of amino acid homorepeats in E-HRPs and NonE-HRPs.** (**A**) The proportion of different amino acid HR types in essential and non-essential human HRPs with structural information in the Protein Data Bank (PDB). n represents the total number of HRs of a particular amino acid type in the PDB. (**B**) Heatmap representing the proportion of different structural conformations corresponding to the HRs of different amino acid types in essential and non-essential human HRPs. Numbers in each cell represent the number of HR entries of different amino acid types. P-value was computed using Fisher's exact test. Rows that contain both cells with non-zero values were used to compute significance estimates.

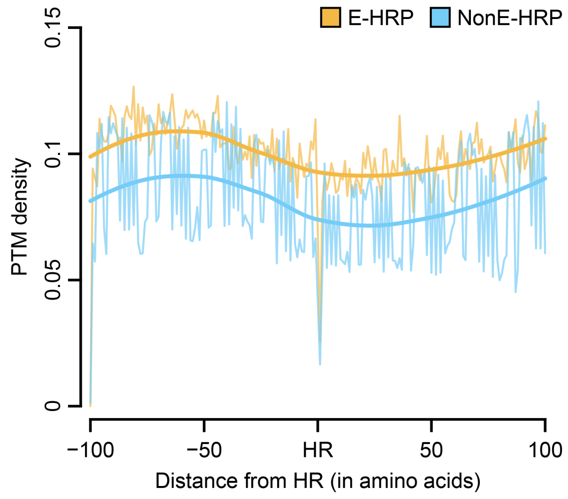


**Fig. S5. Distribution of PTMs across HRs in E-HRPs and NonE-HRPs.** Line plot showing the PTM density around 100 amino acid residues N- and C-terminus to the HR in both classes of HRPs.

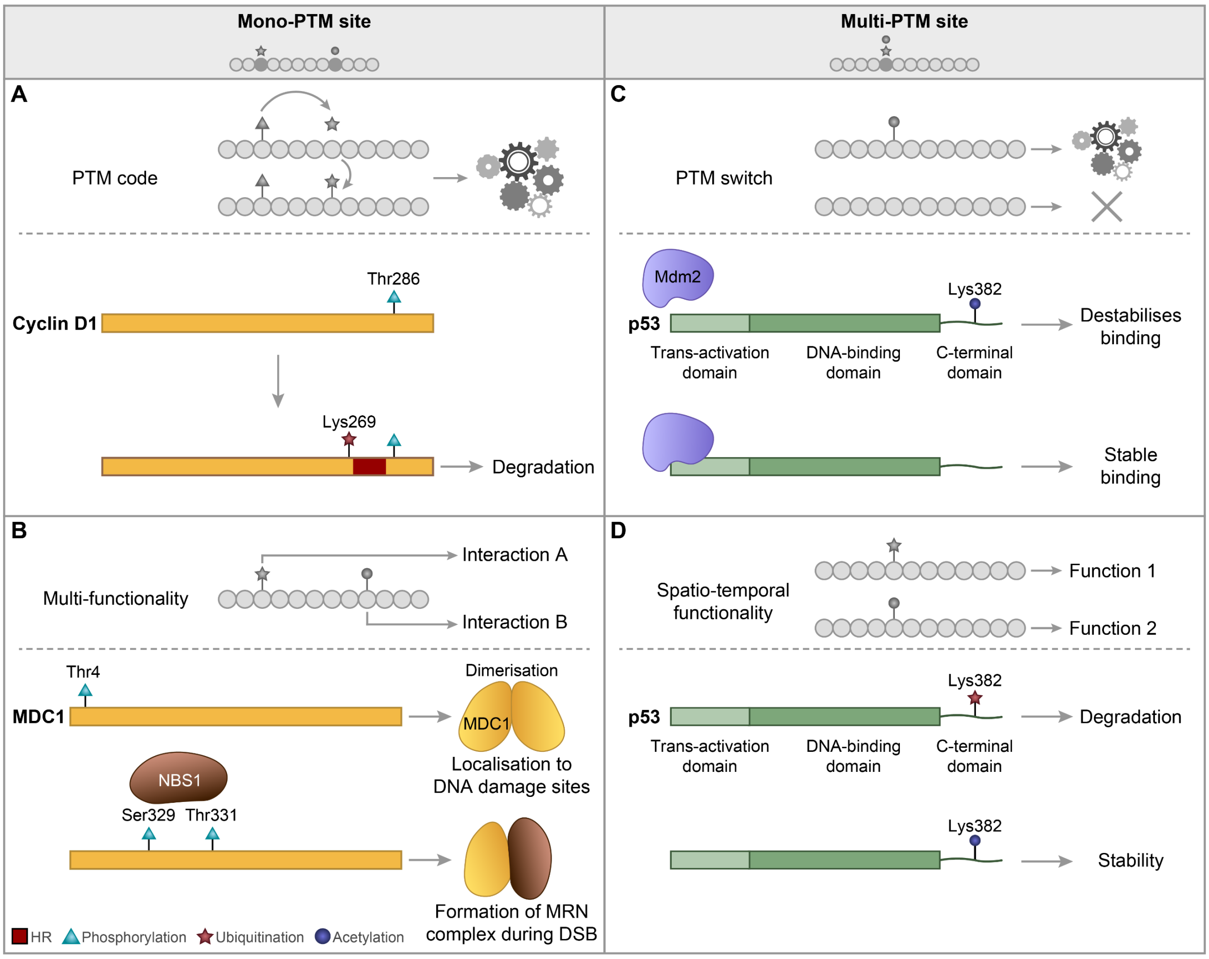


**Fig. S6. Mode of regulation of protein stability/function by mono- and multi-PTM sites in E-HRPs and E-NonHRPs.** (**A**) Phosphorylation at the mono-PTM site Thr286 of E-HRP Cyclin D1 promotes ubiquitination at Lys269 (*1-4*), which consequently leads to the degradation of the protein. (**B**) Phosphorylation at Thr4 of E-HRP MDC1, which is involved in double-stranded break (DSB) repair, allows dimerization of the protein (*5*). This helps MDC1 to localize to DNA damage sites. Phosphorylation at Ser329 and Thr331 facilitates binding to NBS1 (*6*), which then forms the MRN complex during DSB repair. (**C**) Acetylation at Lys382 position of E-NonHRP p53 destabilizes its binding with E3 ubiquitin ligase Mdm2 (*7*), acting as a switch that destabilizes the interaction. (**D**) Ubiquitination at Lys382 position of p53 leads to protein degradation, whereas, acetylation provides stability (*8*), exemplifying the roles of multi-PTM sites.

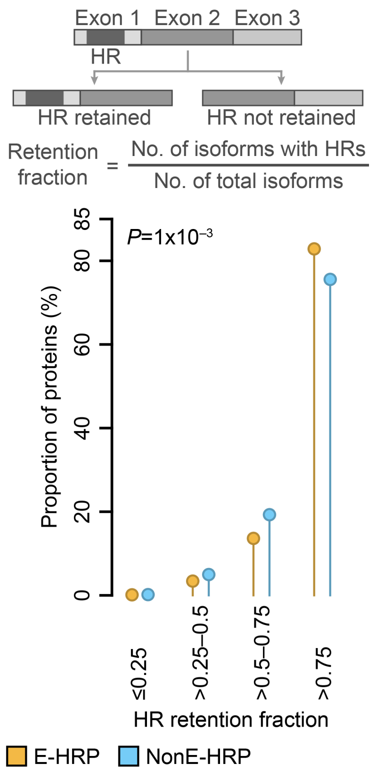


**Fig. S7. Alternatively spliced events among E-HRPs and NonE-HRPs.** Distribution showing the extent of HR retention in E-HRP and NonE-HRP isoforms. Statistical significance was estimated using Fisher's Exact test.

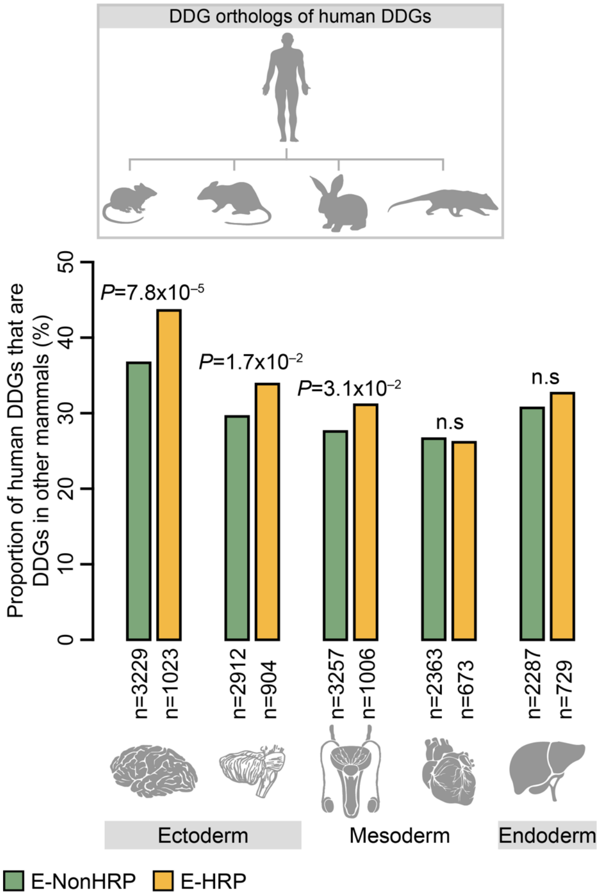


**Fig. S8. Developmental dynamicity of E-HRPs in brain is evolutionarily conserved across mammals.** Proportion of E-HRPs and E-NonHRPs that are developmentally dynamic in different organs in humans and non-human mammals. Statistical significance was assessed using Fisher's Exact test.

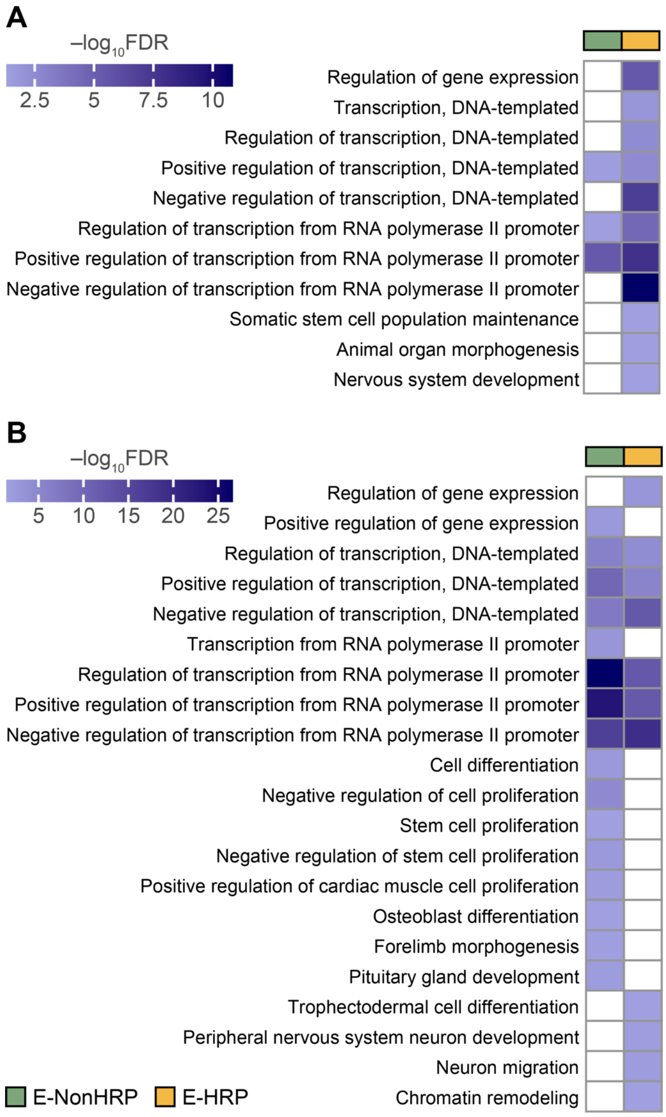


**Fig. S9. HAR-flanked E-HRP and E-HRP transcription factors are involved in embryonic and brain development.** Heatmap showing significantly enriched (FDR<0.05) Gene Ontology biological process (GO-BP) terms among (**A**) E-NonHRPs and E-HRPs and (**B**) E-NonHRPs and E-HRPs transcription factors that are flanked by HARs.

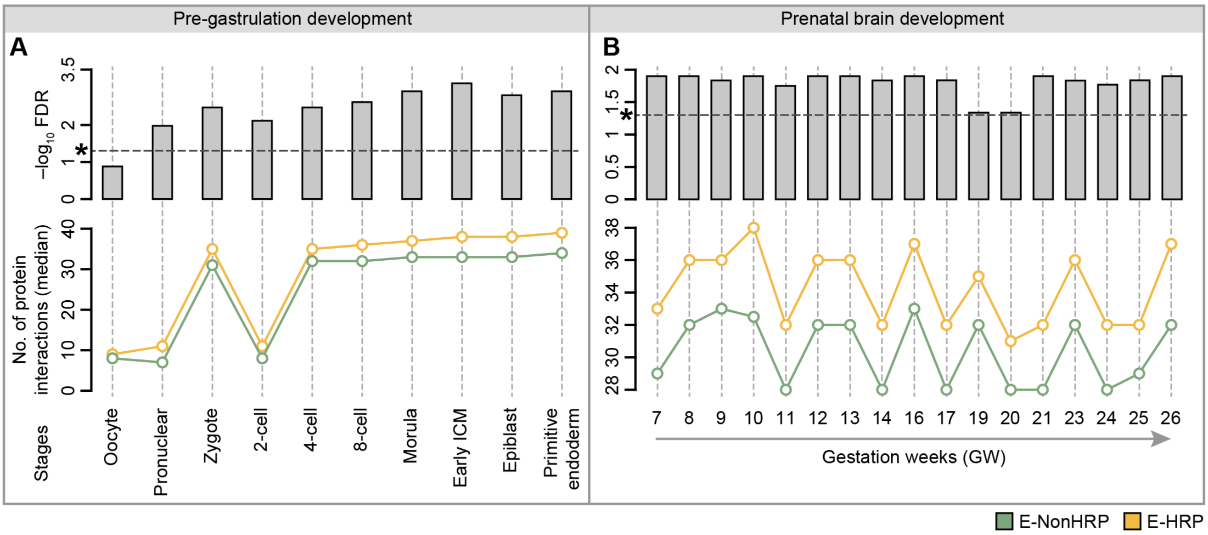


**Fig. S10. E-HRPs show higher number of protein-protein interactions across different stages of development.** Distribution showing the median number of protein interactions of E-HRPs and E-NonHRPs during different stages of (**A**) pre-gastrulation and (**B**) prenatal brain development (bottom panels) and the corresponding significance estimates (top panels). The different developmental stages are given in the X-axes. Correction for multiple testing to estimate significance was done using FDR. In the top panel the horizontal dotted line highlighted with an asterisk denotes the significance line. The FDR values above the dotted line represent statistical significance. ICM stands for Inner cell mass.

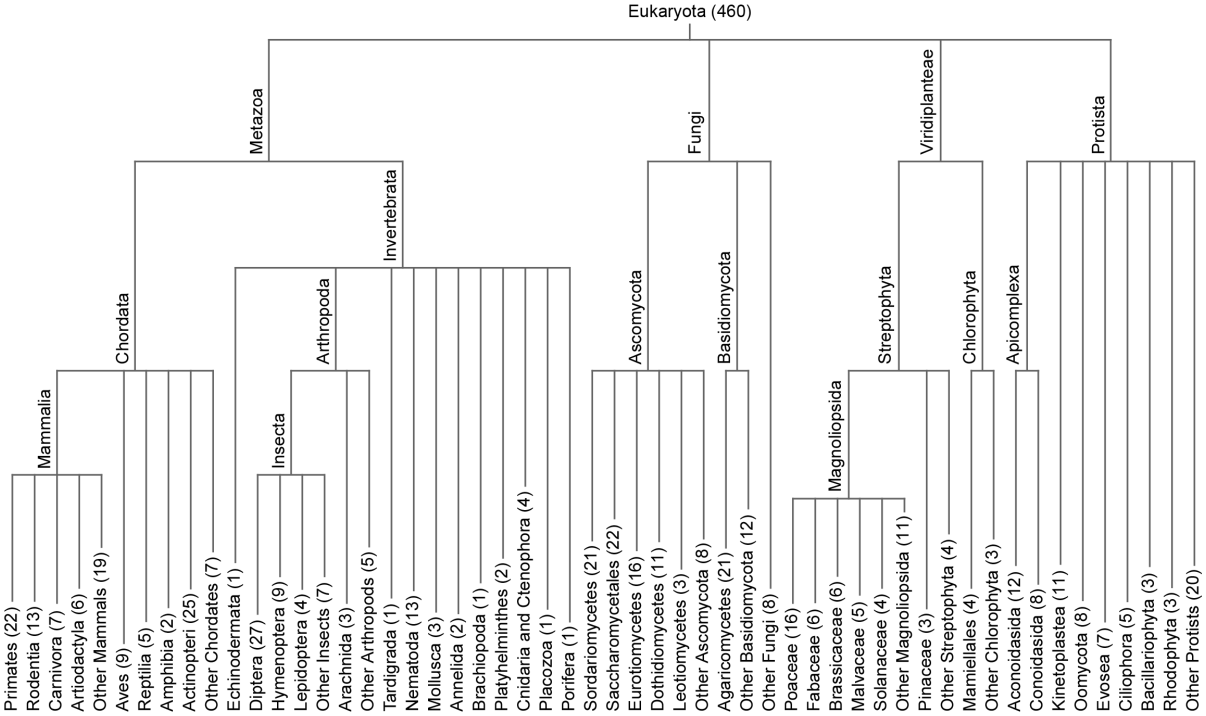


**Fig. S11. Phylogenetic tree depicting the taxonomic classification of the eukaryotic species considered for gene-age estimations.** Number in parenthesis represents the number of species in each taxon. Branch lengths do not correspond to any phylogenetic attributes.

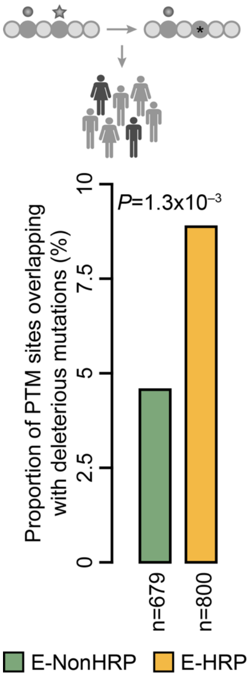


**Fig. S12. Mutations in PTM sites in E-HRPs tend to be more deleterious.** Proportion of PTM sites, mutations in which lead to deleterious effects, in the different classes of proteins. n denotes the number of PTM sites that have non-synonymous substitutions. P-value was computed using Fishers Exact test.

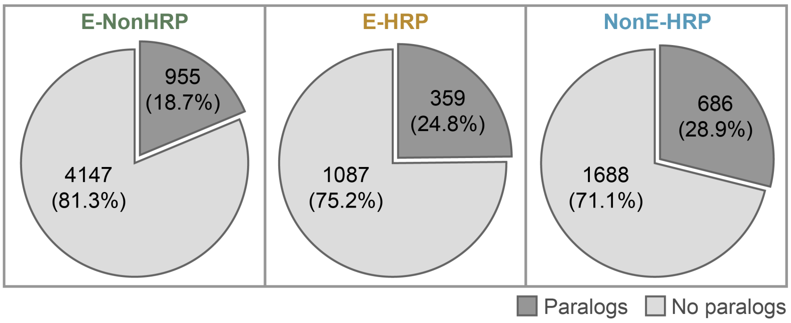


**Fig. S13. Proportion of gene duplicates in different classes of proteins.** The numbers in the pie-chart represent the number of proteins and the proportion of genes with and without paralogs in a class are presented as percentages.

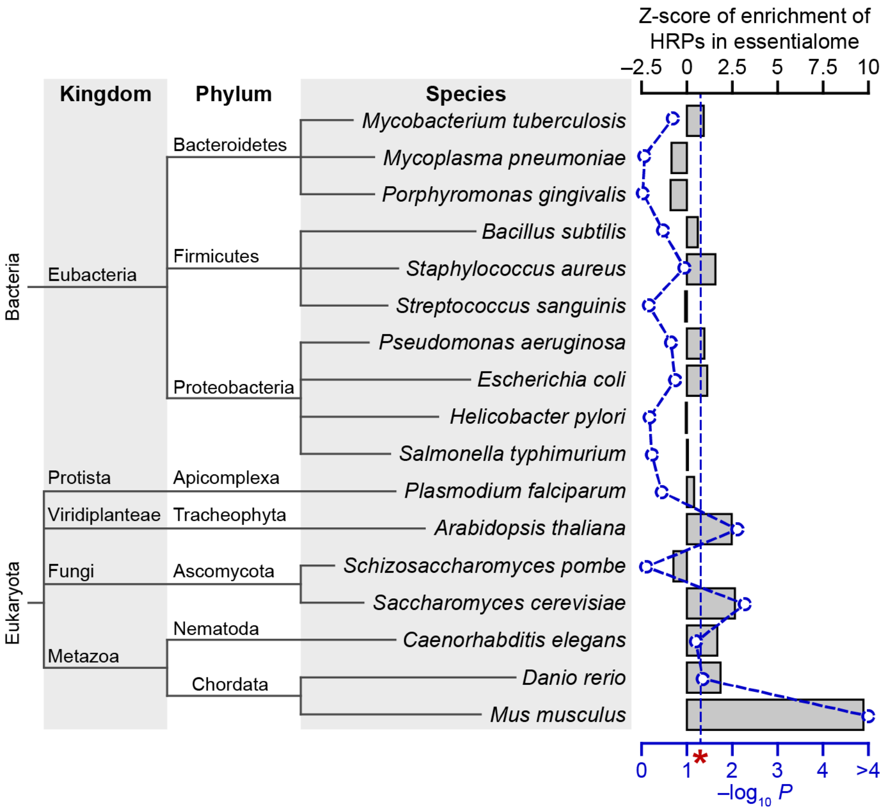


**Fig. S14. Enrichment of proteins with amino acid homorepeats (HRPs) in the essentialomes of 17 species.** The bars provide the Z-scores (top X-axis) corresponding to each species (Y-axis), while the line plot provides the $-\log_{10}$ P-values (bottom X-axis) for enrichment. Z scores and P values were computed using permutation test from 10,000 randomizations, for each species. The vertical dotted straight line denoted by red asterisk represents the $-\log_{10}$ P-values significance cut-off.

**Table S1. Details of studies from which human essentialome was assembled.**

| Description | Number of genes | Cell line(s) | Reference |
|---|---|---|---|
| Extensive mutagenesis in haploid cells to identify genes for viability and fitness | 2498 | HAP1, KBM7 | (*9*) |
| Genome-wide CRISPR/ Cas9 screens and gene-trap method to identify genes required for proliferation and survival | 2709 | KBM7, K562, Jiyoye, Raji | (*10-12*) |
| Genome-wide CRISPR/ Cas9 screen employing negative selection to identify genes affecting fitness negatively | 4048 | A375, DLD1, GBM, HCT116, HELA, RPE1 | (*11-13*) |
| Genome-wide CRISPR/ Cas9 screen of dying cells to identify genes essential for oxidative phosphorylation | 191 | K562 | (*14*) |
| Literature evidence (Review article) | 574 | Across cell lines | (*15*) |
| Analysis of 17 genome-scale knockout screens in human cells using genome-scale gRNA libraries | 682 | Across cell lines | (*16*) |
| Genome-wide loss-of-function screen using CRISPR/ Cas9 affecting normal growth and survival | 1668 | hPSC | (*17*) |
| Genome-scale loss-of-function screen using inducible Cas9 used to identify genes involved in cell fitness | 2347 | hPSC, HAP1 | (*11, 12, 18*) |
| Genome-wide CRISPR/ Cas9 screen to identify genes involved in cell survival and sensitivity to chemotherapeutics | 1058 | PANC-1 | (*11, 12, 19*) |

**Table S2. Compendium of large-scale datasets used in the study.**

| Dataset | Description |
|---|---|
| ***Sequence-related features*** | |
| Human proteome | Obtained from UniProtKB Swiss-Prot (Release_2021_02). The dataset comprised of 20,395 human reviewed protein sequences. |
| Human Eukaryotic Linear Motifs (ELM) | We curated and mapped the ELMs from ELM database (*20*). The analysed dataset contained 1,625 ELMs in 1,218 proteins. |
| Human protein isoforms | Non-canonical sequences of proteins obtained from Swiss-Prot (Release_2021_04). The dataset comprised of 10,530 human reviewed protein sequences with 32,506 total isoforms. |
| Human paralogs | We obtained and curated 2,983 human paralog pairs encompassing 4,485 proteins from Dandage *et al.* (*21*). |
| ***Features related to molecular interactions*** | |
| Human protein-protein interaction network | A comprehensive human experimentally annotated physical protein-protein interaction network was assembled from BioPlex (*22*), CORUM (*23*), DIP (*24*), HINT (*25*), PathwayCommons (*26*), HuRI (*27-32*) and BIOGRID (only physical interactions which are not identified using Yeast2Hybrid system) (*33*), and high-throughput studies (*29*). The curated network consisted of 17,160 proteins and 3,55,718 protein-protein interactions. |
| Human protein-complex network | Human protein-complex network was obtained from hu.MAP 2.0 (*34*). Nearly 15,816 mass spectrometry experiments were used to develop a machine learning framework to predict protein complex assemblies. The curated dataset consisted of 9,940 proteins which formed 6,914 protein complexes. |
| Human phase separated condensates | The dataset was assembled from different databases such as DrLLPS (*35*), LLPSDB (*36*), PhaSepDB (*37*) and PhasePro (*38*). |

| Dataset | Description |
|---|---|
| | The curated dataset contained 3,855 proteins that separate in 65 condensates. These condensates were manually assigned to different biological functions by extensive literature search. |
| Human transcription factor (TF)-gene target regulatory network | We assembled the TF-target network from several databases such as hTFTarget (*39*) and TRRUST v2 (*40*) as well as high-throughput studies (*41, 42*). The curated network consists of 1,115 TFs and 19,026 gene targets and 22,99,409 regulatory interactions. |
| Human RNA Binding Proteins (RBP)-mRNA target regulatory network | The dataset was obtained from RNAct (*43*), and filtered for only experimentally verified interactions and canonical transcript targets. The curated network consisted of 77 RBPs and 3,515 mRNA targets and 17,558 regulatory interactions. |
| Human protein Post-Translational Modifications (PTMs) | We obtained human PTMs from dbPTM (*44*). The curated dataset consisted of a total of 3,56,770 PTMs of 60 different PTM types spanning across 17,829 proteins. |
| ***Features related to structure and protein disorder*** | |
| Human intrinsic protein disorder | We predicted the intrinsic disorder in human proteins using AUCpreD (*45, 46*). Disorder percentage for each protein was computed as the fraction of the total number of disordered residues, over length of the protein. |
| Human homorepeat (HR) structures | We downloaded all human protein structures from Protein Data Bank (*47*). The secondary structure annotations for each structure was obtained from DSSP (*48, 49*). A total of 149 HRs in 145 proteins were identified to have 201 total secondary structures. We compared the number and type(s) of secondary structure(s) adopted by each HR in E-HRPs and NonE-HRPs. |
| ***Regulatory proteins*** | |

| Dataset | Description |
|---|---|
| Human transcription factors (TFs) | List of human transcription factors was collated from experimentally verified high-throughput studies (*41, 42, 50, 51*), and hTFTarget (*39*) and TRRUST v2 (*40*) databases. The curated dataset included 1,608 transcription factors. |
| Human RNA binding proteins (RBPs) | Human RBPs were assembled from experimentally verified high-throughput studies (*52*) and RNAct database (*43*). The curated dataset consisted of 354 RBPs. |
| ***Spatio-temporal expression*** | |
| Human tissue-specific proteomes | Tissue-specific proteomes were obtained from Human Protein Atlas (*53*). Proteins were considered to be expressed in a tissue based on their reliability (either 'Approved', 'Enhanced' or 'Supported') and levels (either 'High', 'Medium' or 'Low'). The curated dataset contained 10,551 proteins across 46 tissues. |
| Organismal developmentally dynamic proteins | Development of seven different organs across five different species including humans, in a temporal manner was studied (*54*). Protein products of genes which show temporal dynamicity in terms of its mRNA expression across different stages of organ development were classified as developmentally dynamic proteins in a given organ for each species studied. |
| Human developmental stage-specific proteomes (pre-implantation) | Pre-gastrulation stage-specific proteomes were obtained from two studies (*55, 56*). We considered genes with normalised averaged FPKM values ≥ 1 across 7 developmental stages from Stirparo *et al.* (*55*) and genes with RPKM value > 0.5 across 7 stages from Madissoon *et al.* (*56*). These two datasets were combined to assemble the pre-gastrulation developmental proteome consisting of 12,702 proteins spread across 10 stages. |

| Dataset | Description |
|---|---|
| Human developmental stage-specific proteomes (brain) | Prenatal brain stage-specific proteomes were assembled from two studies (*57, 58*). Genes expressed between gestation week (GW) 7 and 28 were identified by Fan *et al.* (*57*). In this dataset, genes with normalized counts > 0 in each stage were considered, irrespective of region. Genes were considered only if they were expressed in atleast 5 cells. Data from Pons, in the brain stem, were excluded. Genes expressed in human prefrontal cortex between GW08 and GW26 were identified by Zhong *et al.* (*58*). Genes with TPM values > 1, expressed in atleast 3 cells were considered. These two datasets were combined to assemble the prenatal brain developmental proteome consisting of 16,706 proteins spread across 17 stages. |
| ***Features related to evolution*** | |
| Human protein 1:1 orthologs | The ortholog information was obtained from OMA browser database (*59*) and filtered for 1:1 orthologs. The curated data consists of 18,51,515 orthologs of 17,770 human proteins across 459 eukaryotes. |
| Human accelerated regions (HARs) | We retrieved 3,168 HARs and genes flanking the HARs from Girskis *et al.* (*60*). For identifying HARs active in brain development, we considered the 1,524 HARs that showed activity in the Capture Massively Parallel Reporter Assays (caMPRA) in the neuronal cells. |
| ***Phenotype-related datasets*** | |
| Human essentialome | We collated 6,548 essential genes/proteins from 9 mid- to high-throughput studies (Refer **Table S1**). |
| Human deleterious mutations | About 321,643 human PTMs pertaining to 59 types were mapped to about 4 million non-synonymous single nucleotide variations |

| Dataset | Description |
|---|---|
| | (nsSNVs). Using a relative pathogenicity score (rps) cut-off > 95, about 135 sites across 32 diseases were predicted to be have the PTM site-disease associations (*61*). We have mapped these deleterious PTM site mutations to E-HRPs and E-NonHRPs for studying the differences in distribution of deleterious disease associated PTM-site? mutations across the two classes of essential proteins. |
| ***Datasets of other species*** | |
| Mouse protein PTMs | We retrieved and curated a set of 64,208 PTMs in 32,104 mouse proteins from dbPTM (*44*). |
| Essentialomes and proteomes of other species | We obtained the list of essential genes/proteins across different species from Database of Essential Genes (DEG15) (*11*), which collates essentialome information from various low-, mid- and high-throughput experimental studies. Species with atleast 50 essential genes were considered for the study. We selected 10 representative bacterial species spanning 3 different phyla, with the most number of essential genes in each of the phyla.<br><br>Proteome of *Saccharomyces cerevisiae* was obtained from SGD (*62*), while that of all other species was obtained from UniProt Reference Proteomes database. |

**Table S3. Conditional probabilities of high interactability of E-HRPs across diverse networks.**

| Conditional probability | PPI degree | PPI link communities | TF-target outdegree | RBP-RNA outdegree |
|---|---|---|---|---|
| P (High interactability \| E-HRP) | 0.53 | 0.51 | 0.43 | 0.46 |
| P (E-HRP \| High interactability) | 0.13 | 0.13 | 0.28 | 0.42 |

We classified the number of interactors/targets of all proteins/TFs/RBPs into tertile bins, and defined the proteins/TFs/RBPs that fall in the bin with the highest interactions as those with high interactability.

**Table S4. Taxonomic details of species considered for estimating sequence identities.**

| Species code | Species name | Taxon |
|---|---|---|
| PANTR | *Pan troglodytes* | Primates |
| GORGO | *Gorilla gorilla gorilla* | Primates |
| MACNE | *Macaca nemestrina* | Primates |
| RATNO | *Rattus norvegicus* | Mammals |
| MOUSE | *Mus musculus* | Mammals |
| CANLF | *Canis lupus familiaris* | Mammals |
| PIGXX | *Sus scrofa* | Mammals |
| HORSE | *Equus caballus* | Mammals |
| CHICK | *Gallus gallus* | Chordate |
| CHRPI | *Chrysemys picta bellii* | Chordate |
| XENTR | *Xenopus tropicalis* | Chordate |
| LEPOC | *Lepisosteus oculatus* | Chordate |
| LATCH | *Latimeria chalumnae* | Chordate |
| STRPU | *Strongylocentrotus purpuratus* | Metazoa |
| DROME | *Drosophila melanogaster* | Metazoa |
| CAEEL | *Caenorhabditis elegans* | Metazoa |
| LOTGI | *Lottia gigantea* | Metazoa |
| CAPTE | *Capitella teleta* | Metazoa |
| HYPAI | *Trichoderma atroviride* IMI 206040 | Fungi |
| CANAL | *Candida albicans* SC5314 | Fungi |
| YEAST | *Saccharomyces cerevisiae* S288C | Fungi |
| SCHPO | *Schizosaccharomyces pombe* 972h- | Fungi |
| MUCCI | *Mucor circinelloides* | Fungi |

| **Species code** | **Species name** | **Taxon** |
|---|---|---|
| ORYSJ | *Oryza sativa* Japonica Group | Viridiplantae |
| ARATH | *Arabidopsis thaliana* | Viridiplantae |
| THECC | *Theobroma cacao* | Viridiplantae |
| KLEFL | *Klebsormidium flaccidum* | Viridiplantae |
| CHLRE | *Chlamydomonas reinhardtii* | Viridiplantae |
| PLAF7 | *Plasmodium falciparum* 3D7 | Protista |
| TOXGV | *Toxoplasma gondii* VEG | Protista |
| PHYIT | *Phytophthora infestans* T30-4 | Protista |
| DICDI | *Dictyostelium discoideum* | Protista |
| CAPO3 | *Capsaspora owczarzaki* ATCC 30864 | Protista |
